# Genomic insights into photosymbiosis in giant clams: Comparisons with coral strategies

**DOI:** 10.1101/2025.11.20.689417

**Authors:** Taiga Uchida, Hiroshi Yamashita, Go Shimada, Mayumi Kawamitsu, Eiichi Shoguchi, Chuya Shinzato

**Author notes:** Corresponding Author: Chuya Shinzato.

## Abstract

Giant clams are representative bivalves in coral reef ecosystems that host photosynthetic dinoflagellates extracellularly and rely on their photosynthates, functioning as “solar-powered animals.” Unlike corals, which harbor intracellular dinoflagellates, molecular mechanisms and evolutionary history underlying this symbiosis remain largely unknown. Here, we integrated chromosome-scale genome assembly, transcriptome profiling, and bleaching experiments involving *Tridacna crocea* to explore the genetic basis of extracellular symbiosis. Signals associated with sterol transport by Niemann-Pick disease type C2 (NPC2) transporters and carbon-concentrating mechanisms suggest that giant clams share some nutrient exchange strategies with corals. Strikingly, the nitrate transporter NRT2, a “plant-like” gene previously thought to be absent in animals, represents an unexpected evolutionary retention that enables nitrate-based nutrient supply, highlighting a fundamental divergence from coral symbiosis. Together, our findings reveal both conserved and distinct molecular strategies of photosymbiosis in reef-dwelling marine invertebrates and provide insights into evolution and ecological resilience of coral reef ecosystems.

## Introduction

Symbiosis is a widespread feature across the tree of life and has been pivotal in acquisition of novel traits and adaptations throughout evolutionary history. Among various forms of symbiosis, “photosymbiosis” refers to associations between heterotrophic hosts and phototrophic microbes, such as cyanobacteria or unicellular algae. In animals, photosymbiosis has evolved independently in multiple lineages, including Porifera, Cnidaria, Xenacoelomorpha, Platyhelminthes, Mollusca, Tunicata, and even Vertebrata (*1*). In general, symbionts provide photosynthates to their hosts, while hosts offer inorganic nutrients, such as carbon, nitrogen, and phosphorus, and a protective microhabitat in return (*2*).

Intracellular symbiosis between reef-building scleractinian corals and dinoflagellates of the family Symbiodiniaceae (zooxanthella) represents an example of photosymbiosis, which underlies biodiversity in coral reef ecosystems. However, this symbiotic relationship is fragile and can collapse under thermal stress, resulting in a phenomenon known as coral bleaching. With increasing anthropogenic challenges such as global warming, coral bleaching has become an urgent global issue (*3*, *4*). In response, there has been a surge in research focusing on molecular mechanisms underlying coral-dinoflagellate symbiosis. Since the first report of the coral and symbiotic dinoflagellate draft genome assembly in 2011 and 2013 (*5*, *6*), genomic and transcriptomic studies have significantly advanced our understanding of the molecular basis of photosymbiosis in corals and related cnidarians (reviewed by Shinzato and Yoshioka (*7*)). Dinoflagellates also engage in symbiosis with a variety of non-cnidarian hosts, including molluscs (bivalves and nudibranchs), xenacoelomorphs, and sponges (*8*, *9*). Nevertheless, molecular mechanisms underlying these associations remain poorly understood, largely due to the lack of genomic resources for these host organisms.

Giant clams (genera *Tridacna* and *Hippopus*, subfamily Tridacninae, Family Cardiidae) are conspicuous tropical bivalves of Indo-West Pacific coral reef ecosystems, well known for their extremely large size and colorful mantles (*8*) (Fig. 1A). They engage in symbiotic relationships with algae of the Family Symbiodiniaceae. These symbionts reside extracellularly, not intracellularly, in a highly branched tubular structure in the host outer mantle, called the “zooxanthellal tube system” (*10*). Giant clams receive photosynthetic products from dinoflagellates, in addition to filter feeding like most bivalves. (*11*, *12*). Clams discharge intact and photosynthetically active dinoflagellate cells in their fecal pellets, which are capable of colonizing and proliferating in coral larvae, suggesting a potential source of symbiotic algae for corals (*13–16*). In recent years, giant clams have gained attention as models for photosymbiosis research and as targets for aquaculture and conservation (Fig. 1A), leading to rapid advances in genomic studies. The Aquatic Symbiosis Genomics project led by the Wellcome Sanger Institute (*17*), along with multiple independent research teams, has contributed to a surge in genome sequencing efforts. Between December 2023 and June 2025, chromosome-scale genome assemblies of five *Tridacna* species were reported (*18–23*) (including preprints, with one species not yet publicly released). Furthermore, genomes of three *Fragum* species (Subfamily Fraginae, Family Cardiidae), another major lineage that engages in symbiosis with dinoflagellates, were reported in 2024 by the same project (*24–26*). These advances mark the beginning of a new era in which bivalve photosymbiosis can be systematically investigated through the lens of genomics.

**Fig. 1.**
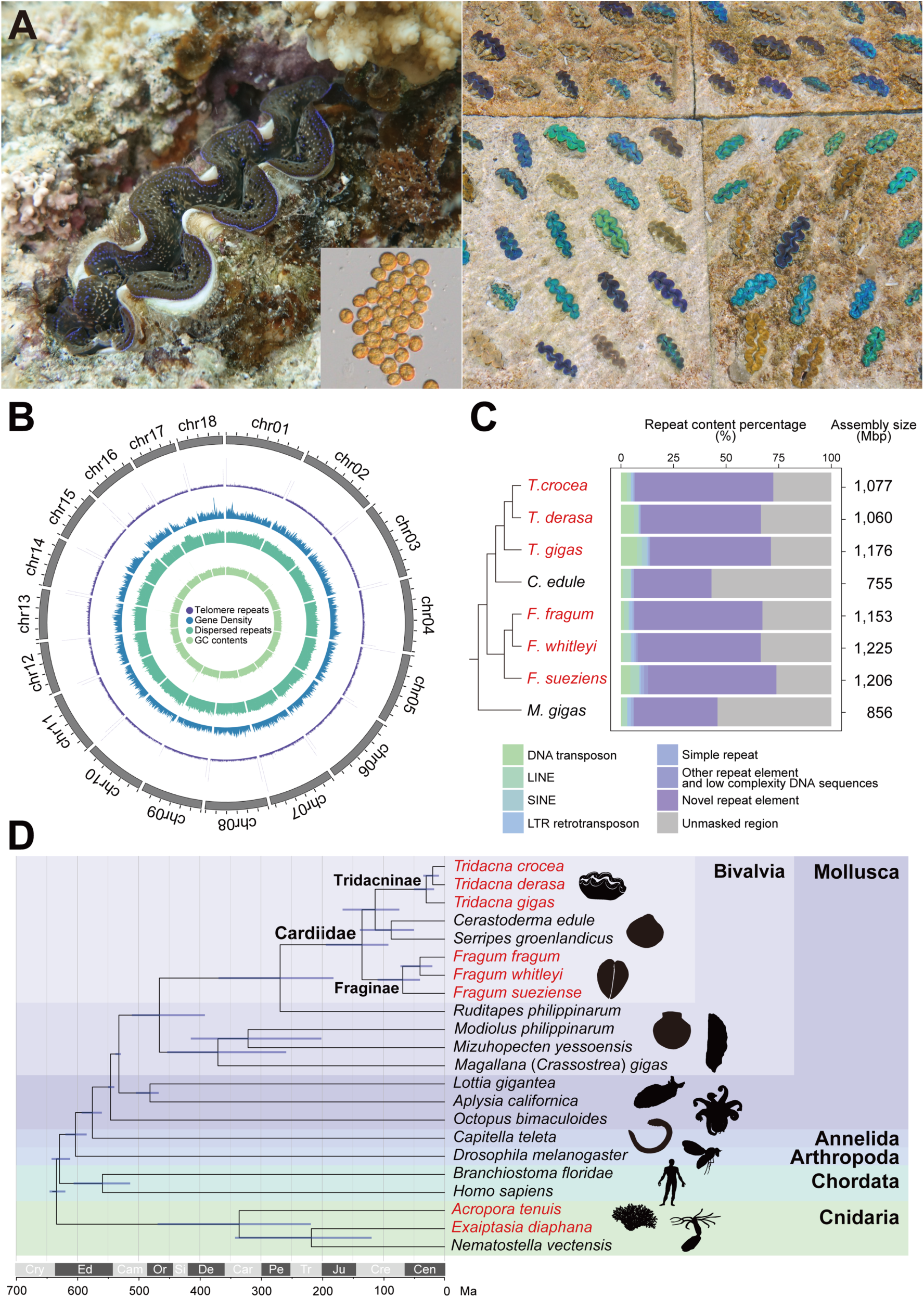
Overview of the *Tridacna crocea* genome and phylogenetic placement. (**A**) *Tridacna crocea* embedded in coral rock at a shallow depth on the coral reef of Ishigaki Island, Okinawa, Japan, and symbiotic algae (left) and those reared at the Miyakojima City Center for Stock Enhancement, located on Miyako Island (Miyakojima, Okinawa, Japan) (right). (**B**) Genomic landscape of *T. crocea* genome haplotype 1, showing telomere repeats (AACCCT)n (purple; 0-2,000 units) counted for 100-kb non-overlapping windows, protein-coding gene density (blue; 0-100 genes/Mb), interspersed repeat density (green; 0-100%), and GC content (light green; 30-50%). Gene density, interspersed repeat density, and GC content were calculated for 1-Mb overlapping windows with 100-kb step size. A Circos plot was created using interacCircos v.1.14.0 (*131*). * > 300 telomere repeats in the terminal 10-kbp region. (**C**) Genome sizes (right) and repeat contents (middle) of chromosome-scale assembled genomes of bivalves. The cladogram represents the phylogenetic relationship of bivalve species estimated in the present study (left). Only chromosome-scale genomes are compared. The “Novel” repeat class represents elements without any BLASTN or BLASTX hits in repeat databases. (**D**) Phylogenetic relationships and divergence times of 22 metazoan species. A maximum likelihood molecular phylogenetic tree was constructed using 95,883 amino acid residues from 402 single-copy orthologs shared by all species. Divergence times (million years ago; Ma) were estimated using six fossil calibrations. Horizontal bars indicate 95% credible intervals of estimated divergence times. Cry, Cryogenian; Ed, Ediacaran; Cam, Cambrian; Or, Ordovician; Sl, Silurian; De, Devonian; Car, Carboniferous; Pe, Permian; Tr, Triassic; Ju, Jurassic; Cre, Cretaceous; Cen, Cenozoic.

Here, we present a chromosome-scale genome assembly of *Tridacna crocea* that ranks among the highest quality assemblies reported for giant clams to date. *T. crocea*, the smallest species in the Subfamily Tridacninae, is widely cultivated and offers practical advantages as an experimental model. Leveraging this new genomic resource with recently published genomes of closely related species, we performed a comprehensive comparative genomic analysis to identify lineage-specific genetic features potentially linked to photosymbiosis. We further conducted tissue-specific transcriptomic profiling in *T. crocea*, with a particular focus on the outer mantle, a specialized structure that houses algal symbionts. In addition, we investigated transcriptomic responses of *T. crocea* to dark-induced bleaching, the collapse of the symbiotic relationship between clams and algal symbionts (*27*, *28*). By integrating genomic and transcriptomic data, we screened for candidate symbiosis-related genes and investigated the evolutionary origins of particularly intriguing candidates. Our findings provide novel insights into the genetic basis of photosymbiosis in giant clams with reference to cnidarians, including corals. These discoveries contribute to a broader understanding of how marine invertebrates maintain symbioses with algal partners.

## Results and Discussion

### Chromosome-Scale Genome Assembly Reveals a High Proportion of Repeat Elements in Photosymbiotic Bivalves

To minimize contamination from algal symbionts, genomic DNA was extracted from gonads of *Tridacna crocea* and sequenced using HiFi and Hi-C technologies. The final assemblies comprised 60 and 56 scaffolds for haplotypes 1 and 2, respectively, with total lengths of 1,077 Mbp and 1,057 Mbp and scaffold N50s of ~61 Mbp. Among them, 18 scaffolds for each haplotype seemingly correspond to chromosome-scale sequences, consistent with the karyotype of 2n = 36 observed in embryos (Fig. S1). These metrics are comparable to other chromosome-scale genomes of tridacnid clams (Table 1 and S1). In the following sections, we report results based primarily on haplotype 1.

**Table 1.**
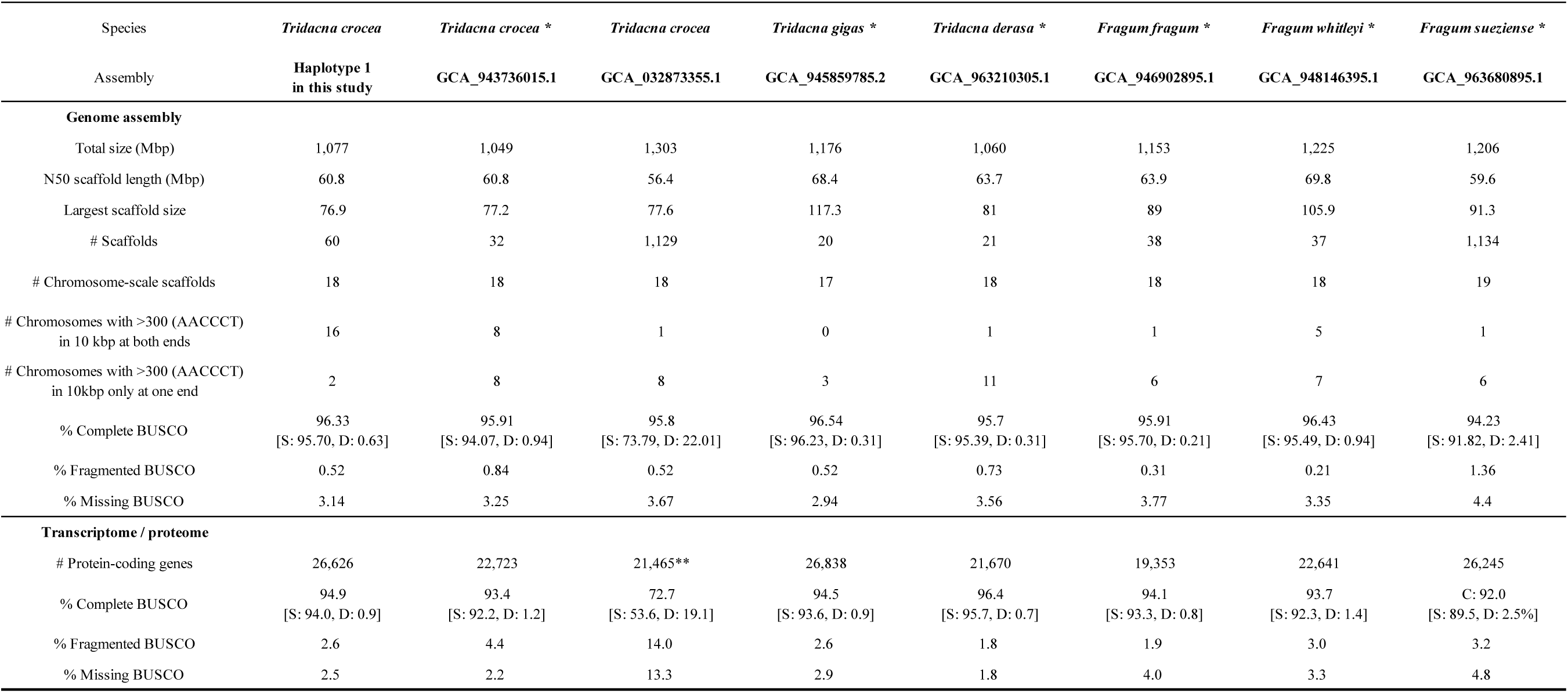
Genome assembly and gene prediction statistics. All gene models except those for GCA_032873355.1 were predicted in this study. BUSCO completeness of the genome and proteome was calculated using compleasm v.0.2.6 and BUSCO v.5.4.7, respectively. S, single-copy BUSCO; D, duplicated BUSCO. Telomere repeats (AACCCT)n were counted with tidk. * RefSeq representative genome (as of 2025/10/1). ** Gene models were downloaded from Li (2023). Chromosome-level genome assembly and annotation of rare and endangered tropical bivalve, *Tridacna crocea.* figshare. Dataset. https://doi.org/10.6084/m9.figshare.24264646

Completeness of the genome evaluated by BUSCO analysis was comparable to those of previously published chromosome-scale genomes of giant clams (Table 1 and S1). To evaluate telomeric completeness, we counted the number of telomeric repeats (AACCCT)n in the terminal 10-kbp regions of each chromosome-scale scaffold. 16 of the 18 chromosomes contained more than 300 telomeric repeats at both ends, while the remaining two chromosomes (chr10 and chr13) had over 300 repeats at one end (Fig.1B and Table 1 and S1), indicating nearly telomere-to-telomere assembly. These values substantially exceed those reported in previously published giant clam genomes, highlighting the contiguity and completeness of our assembly.

We constructed gene models of *T. crocea* and eight species from the Family Cardiidae, whose genome assemblies were publicly available. 26,626 genes were predicted and 94.9% of highly conserved single-copy orthologs in metazoan species were recovered (94.0% single-copy and 0.9% duplicated) for *T. crocea* (Table 1 and S1).

In the process of gene prediction, DNA sequences for repetitive elements (interspersed repeats and low-complexity DNA sequences) in the genomes were identified. For species of *Tridacna*, approximately 67-72% of the genome consisted of repetitive elements (Fig. 1C), which is comparable to *T. maxima* (*23*). Similarly, approximately 66-74% of genomes of species in the Genus *Fragum*, another lineage of photosymbiotic bivalves, consisted of repetitive elements. On the other hand, only 43 and 46% of *Cerastoderma edule* and *Magallana gigas* (*Crassostrea gigas*), non-photosymbiotic bivalves, were repetitive elements (Fig. 1C). Assembly sizes of *Tridacna* and *Fragum* species were approximately 1,060-1,225 Mbp, while those of *C. edule* and *M. gigas* were 755 and 856 Mbp, respectively (Fig. 1C). These results indicate that repetitive elements expanded during evolution of two lineages of photosymbiotic bivalves, resulting in larger genome sizes compared to non-photosymbiotic relatives. According to repeat-content estimates provided for 58 representative bivalve genomes by Chen *et al.* (*29*), the median proportion was 45.6% (interquartile range: 37.7–52.1%). Only four species (6.9%), including *T. crocea* and *Conchocele bisecta*, which form symbiotic relationships with chemosynthetic bacteria (*30*), exceeded 65%. This indicates that the repeat content of *Tridacna* and *Fragum* is exceptionally high among bivalves.

Li *et al.* (*23*) assigned a more detailed annotation to repetitive elements in the *T. maxima genome.* Some types of TEs (LINEs, LTRs, and DNA transposons) were estimated to have undergone rapid expansion during or shortly after the origin of photosymbiosis and two complementary perspectives were proposed. On the one hand, photosymbiosis may have facilitated expansion of transposable elements through processes such as horizontal transfer or relaxed immune suppression. On the other hand, expansion of TEs could have acted as a driving force for emergence of this new adaptive trait by providing novel regulatory elements and gene duplications (*31–33*). Indeed, a similar pattern of repetitive contents in host genomes is observed in the zooxanthellate cnidarian, *Palythoa* (*34*), in addition to *Tridacna*, *Fragum* (this study), and a chemosynthetic bivalve, *Conchocele bisecta* (*30*). Our findings from a comprehensive comparison of bivalve genomes support a possible relationship between symbiosis and repetitive element expansion.

### Phylogenetic Placements and Divergence Time Estimates Provide Clues to the Evolutionary History of Photosymbiotic Bivalves

Molecular phylogenetic analysis using amino acid sequences of 402 single-copy orthologs supports the monophyly of genera *Tridacna* (Tridacninae) and *Fragum* (Fraginae) (Fig. 1D). *Cerastoderma edule* (Lymnocardiinae) and *Serripes groenlandicus* (Clinocardiinae) formed a monophyletic group. This group was a sister clade to the genus *Tridacna*. The Genus *Fragum* was placed as a sister clade to this group (Fig. 1D). All phylogenetic relationships in the Family Cardiidae were supported with 100% bootstrap values (Fig. S2). Previous analyses supported the monophyly of Tridacninae and Fraginae, but yielded inconsistent results regarding their relationships (*35–37*). Recent multi-gene phylogenetic analyses have suggested two alternative topologies: (1) Tridacninae and Fraginae form a monophyletic clade, with Lymnocardiinae as the sister group to all other Cardiidae, and (2) Fraginae is placed as the sister group to a clade composed of Tridacninae and Lymnocardiinae (*38*). In both models, symbiotic and non-symbiotic species in the Fraginae were each recovered as monophyletic. Our present results are consistent with the latter topology. Moreover, a recent phylogenomic analysis of representative molluscan relationships, which utilized BUSCO genes from 77 genomes, also recovered a topology consistent with our results (*29*). Taken together, these findings support the hypothesis that photosymbiosis evolved in the common ancestor of the Tridacninae after its divergence from the Lymnocardiinae, and independently in a single lineage within the Fraginae.

Two lineages of photosymbiotic bivalves, *Tridacna* and *Fragum*, were estimated to have diverged ~134.9 million years ago (Ma) (95% credible interval: 91.7-194.0 Ma). *Tridacna* and *Cerastoderma* were estimated to have diverged ~113.3 Ma (95% CI: 73.9-166.5 Ma). The photosynthetic symbiotic relationship seems to have been acquired since then (*9*). These estimates are roughly consistent with those inferred from a recent phylogenomic study of molluscan relationships using BUSCO genes (*29*). Diversification of crown *Tridacna* and *Fragum* occurred approximately 30.1 million years ago (Ma) (95% CI: 17.0-49.9 Ma) and 68.3 Ma (95% CI: 40.1-109.4 Ma), respectively (Fig. 1D). The genus *Acropora*, the most diverse and abundant scleractinian coral in coral reefs, is assumed to have diversified 25–50 Ma, possibly facilitated by global cooling following the Paleocene–Eocene Thermal Maximum (55.8 Ma) and the subsequent Early Eocene Climatic Optimum (51–53 Ma) (*39–42*). In addition, global reef sites are thought to have expanded dramatically during a period between 40 and 20 Ma (*43*). Diversification of *Tridacna* coincides with the timing of *Acropora* diversification and global reef site expansion, suggesting that radiation of giant clams was driven by expansion and ecological diversification of reef habitats.

### Possible Symbiosis-related Genes Highly Expressed in the Outer Mantle

A tissue-specific transcriptomic analysis was performed using RNA-seq data obtained from six tissues of *T. crocea*: adductor muscle, gill (ctenidia), hepatopancreas, kidney (organ of Bojanus), outer mantle, and pedal mantle (Fig. 2A). Principal component analysis (PCA) based on transcripts per million (TPM) values revealed distinct transcriptional profiles of adductor muscle, gill, hepatopancreas, kidney, and mantle tissues, whereas outer and pedal mantle exhibited similar expression patterns (Fig. 2B).

**Fig. 2.**
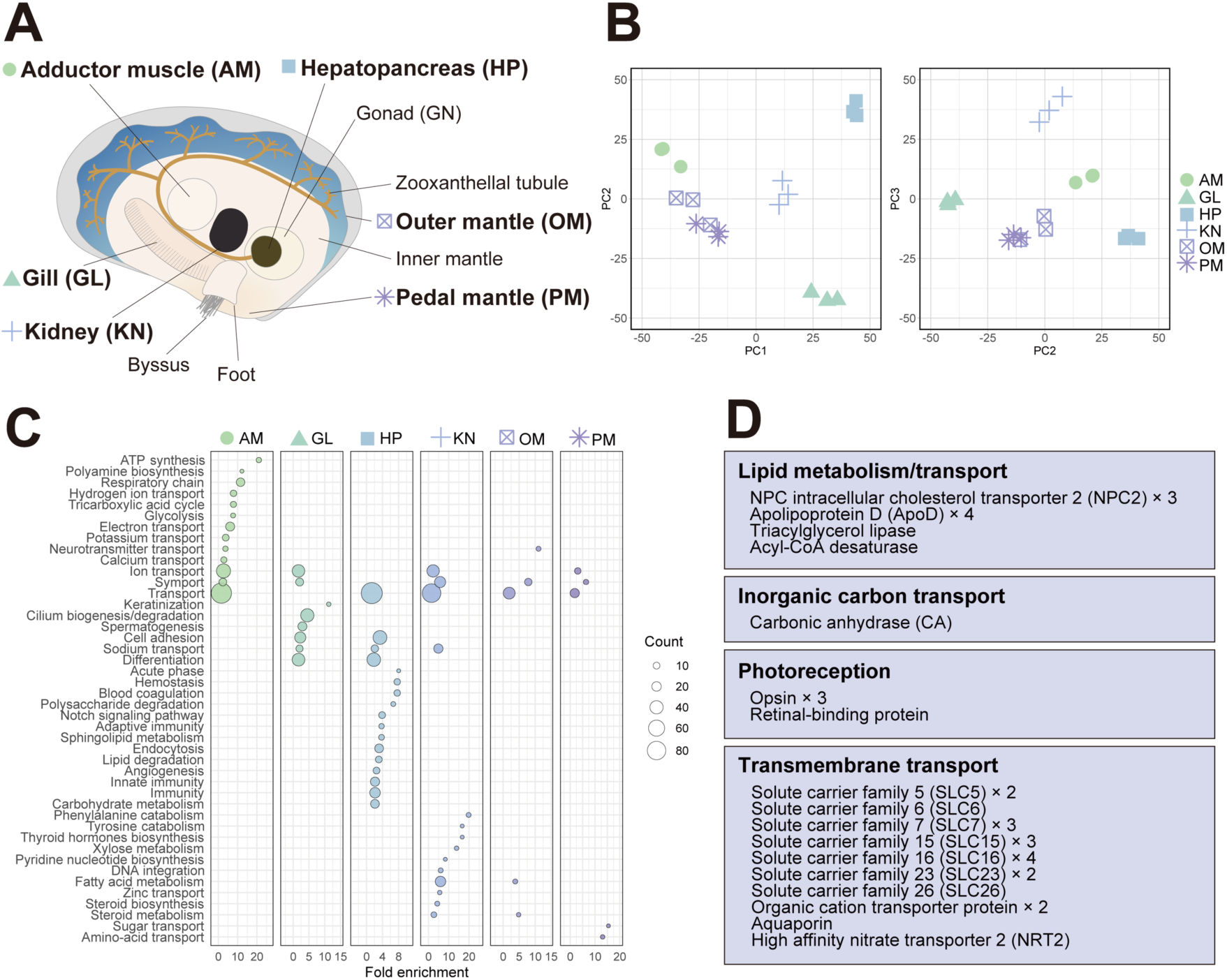
Overview of tissue-specific gene expression patterns of *T. crocea* tissues. (**A**) Schematic diagram of anatomy of *T. crocea*. (**B**) Principal component analysis (PCA) plot of gene expression levels of each sample. Analysis was performed using log-transformed TPM values derived from RNA-seq data by prcomp function in R software. (**C**) Functional enrichment analysis of tissue-specific HEGs based on gene UniProt (UP) terms. Terms of biological function with *p*-values < 0.05 are shown. Results based on molecular function, cellular component, and gene ontology (GO) terms are shown in Table S7-12. (**D**) Manually selected outer mantle-specific HEGs that may be involved in the maintenance of symbiosis with dinoflagellates. AM, adductor muscle; GL, gill; HP, hepatopancreas; KN, Kidney; OM, outer mantle; PM, pedal mantle.

In order to identify genes that characterize each tissue, we examined tissue-specific, highly expressed genes (HEGs): adductor muscle (686), gill (1404), hepatopancreas (1,258), kidney (640), outer mantle (221), pedal mantle (184) (FDR < 0.05 and log_2_ fold change > 1, Fig. S3 and Table S6). Functional enrichment analysis of HEGs revealed transcriptional features of each tissue (Fig. 2C and Table S7-S12). We focused on HEGs specific to the outer mantle, the tissue that harbors symbiotic algae, because some of them may be essential in symbiosis through interactions with symbionts. Outer mantle was characterized by UniProt (UP) keywords and GO terms related to sterol metabolism, fatty acid metabolism, and transport. Multiple copies of genes encoding NPC intracellular cholesterol transporter 2 (NPC2) and apolipoprotein D (ApoD) contributed to enrichment of the UP keywords, *sterol metabolism* and *transport* in the outer mantle. NPC2 is discussed below in detail, and ApoD is discussed in the Supplementary Text and Fig. S4.

Possible symbiosis-related genes were also explored manually from outer mantle-specific HEGs (Fig. 2D). In addition to genes related to lipid metabolism, e.g., NPC2, ApoD, triacylglycerol lipase, and acyl-CoA desaturase, these HEGs included genes involved in photoreception (opsin and retinal-binding protein), and inorganic carbon supply (carbonic anhydrase 2; CA2). In scleractinian corals, CA acts with V-type H^+^-ATPase to form a host carbon-concentrating mechanism (CCM) that enhances algal photosynthesis (*44*, *45*) (Fig. 6). A comparable CCM has been proposed for the giant clam outer mantle, distinct from the inorganic carbon supply for shell formation in the inner mantle, where tubular epithelial cells elevate the CO_2_ concentration around extracellular symbionts (*46–48*). The outer-mantle-specific CA detected here is a plausible component of this system (Fig. 6, see also Supplementary Texts and Fig. S5). Outer mantle-specific HEGs also included many genes involved in transmembrane transport, e.g., solute carrier (SLC) proteins and high affinity nitrate transporter 2 (NRT2). These proteins may be involved in exchange of substances between the host and symbionts. Indeed, several transporter genes have been suggested as candidate genes related to symbiosis between corals and dinoflagellates, due to their upregulation during establishment of symbiosis (*49*, *50*).

### Transcriptomic Responses to the Collapse of Symbiosis

Collapse of the symbiotic relationship between *T. crocea* and algal symbionts was induced by exposure to continuous darkness for two or three months (dark-induced bleaching; Fig. 3A) (*28*). Compared to the control group, the bleached group showed a decrease of approximately 52% in symbiotic algal cells at 2 months and approximately 96% at 3 months (Fig. 3A) (*28*). A transcriptomic analysis was then performed using 3’mRNA-seq data obtained from six tissues of *T. crocea* (adductor muscle, gill, hepatopancreas, kidney, outer mantle, and gonad) (Fig. 2A and S6). In each tissue from bleached clams, 24 to 437 genes were differentially expressed at all time points (Fig. 3B and Table S13). In all tissues, excluding the kidney, there were more genes with decreased expression than those with increased expression. Functional enrichment analysis showed that in gills, which are thought to be involved in nutrient uptake, genes related to transport were downregulated at both time points (Fig. 3C and Table S14, S15). Two of them encode a bicarbonate (HCO_3_^−^) transporter of the SLC4 family (Fig. S10), and one of them was also a gill-specific HEG. The SLC4 transporter in gill epithelium is proposed to export HCO_3_^−^ into the hemolymph after hydration of absorbed CO_2_, supporting the host CCM and the CO_2_ supply to symbionts (*48*, *51*). A significant decline of SLC4 gene expression during bleaching may reflect a decrease in the need for inorganic carbon supply for photosynthesis by symbionts and for shell formation.

**Fig. 3.**
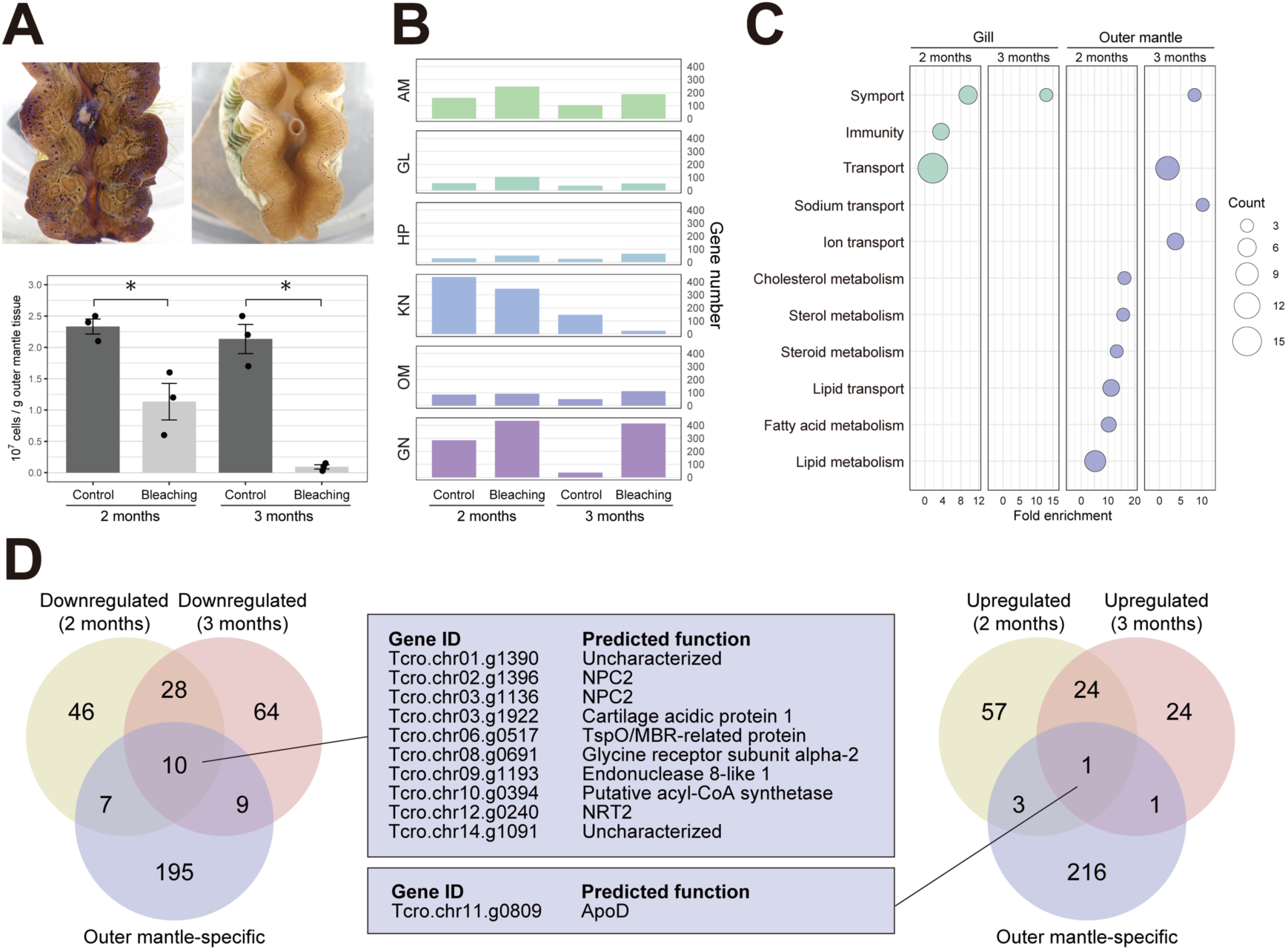
Overview of the dark-induced bleaching experiment and transcriptomic analysis. (**A**) *Tridacna crocea* exposed to natural photoperiod (top left) and continuous darkness (top right) for two months. Dinoflagellate cell density in the outer mantle tissue (bottom). Data were adopted from Uchida *et al.* (*28*) Error bars represent standard errors. * Welch’s *t*-test *p*-value < 0.05. (**B**) Numbers of DEGs in each tissue. (**C**) Functional enrichment analysis of DEGs in gill and outer mantle based on UniProt (UP) keywords. Terms of biological function with *p*-values < 0.05 are shown. Results based on molecular function, cellular component, and gene ontology (GO) terms are shown in Table. S14-17. (**D**) Venn diagram showing numbers of outer mantle-specific HEGs from tissue-specific transcriptomic analysis and DEGs from dark-induced bleaching analysis with predicted functions of genes shared by the three categories assigned by BLASTP searches against the SwissProt database or InterProScan. AM, adductor muscle; GL, gill; HP, hepatopancreas; KN, Kidney; OM, outer mantle; GN, gonad.

In the outer mantle, genes related to lipid metabolism (including sterol and fatty acid metabolism) and lipid transport were downregulated at 2 months (Fig. 3D and Table S16, S17). At 3 months, downregulated DEGs were enriched with genes related to transport, including lipid transport (NPC2), synaptic transmission (acetylcholine receptor subunits), and anion transport (SLC26). Downregulation of genes involved in lipid transport and metabolism may reflect decreased lipid supply from the algal symbiont in this tissue. Downregulation of synaptic transmission genes may reflect a diminished need for responses to light conditions under continuous darkness. Members of the SLC26 family mediate transport of various anions, including chloride (Cl^−^) and other halides, sulfate (SO_4_^2−^), bicarbonate (HCO_3_^−^), oxalate (C_2_O_4_^2−^), and nitrate (NO_3_^−^) (*52*). Expressed outer mantle-specific members and downregulated members of this family may contribute to CCM in this tissue by transporting bicarbonate, like SLC4, or to sulfate or nitrate supply to symbionts, which are essential inorganic nutrients for photosynthetic organisms (discussed below).

Among outer mantle-specific HEGs, 26 showed decreased expression at either 2 or 3 months, whereas only 5 genes exhibited increased expression at these times. Notably, 10 genes showed decreased expression at both 2 and 3 months, while only a single gene displayed increased expression at both time points (Fig. 3D). Among them, two genes encoding NPC2 may contribute to utilization of sterols derived from algal symbionts, whereas NRT2 may be involved in nitrate supply to symbionts, and may have a unique evolutionary history (discussed below in detail).

### Parallel Evolution of Sterol Transport in Animal–Alga Symbiosis: NPC2 at the Host–Symbiont Interface

Because outer mantle-specific HEGs were enriched with lipid metabolism-related genes (Fig. 2C and 2D) and because several of them were downregulated during bleaching (Fig. 3C and D), acquisition of symbiont-derived lipids in the outer mantle is likely important for the host. Indeed, in cnidarians, lipids including triacylglycerols, fatty acids, and sterols (or steryl esters) are thought to be transferred from their symbionts (*2*, *53*, *54*). Also, in Symbiodiniaceae-hosting nudibranchs, it has been proposed that symbiont-derived lipids are transported into host lipid droplets and utilized (*55*). Among these candidates, we focused on NPC2 (Fig. 3D), which is involved in symbiosis between cnidarians and dinoflagellates, by mediating acquisition of symbiont-derived sterols (*54*) (Fig. 6). It is significantly expressed in symbiont-hosting cells or tissues of cnidarians, and upregulated during symbiosis establishment (*50*, *54*, *56–58*).

The *T. crocea* genome includes five probable NPC2 genes, and InterProScan detected MD-2-related lipid-recognition (ML) domain (Pfam: PF02221) in them, as well as in animal NPC2. A molecular phylogenetic analysis was conducted using *T. crocea* NPC2 proteins, some other sequences from the same gene family, and additional cnidarian homologs (Fig. 4A). Two homologous sequences from the single-celled eukaryotic filasterean, *Capsaspora owczarzaki*, were also incorporated following Hambleton *et al.* (*54*). A separate molecular phylogenetic analysis was also performed using a different dataset, focusing on bivalve species relative to *T. crocea* (Fig. 4B). In this analysis, NPC2 sequences from tridacnine clams were classified into six clades based on the tree topology, and were designated TriNPC2.1–6. Among them, TriNPC2.2–6 were strongly supported with bootstrap values ranging from 80% to 100%. Although TriNPC2.1 received low bootstrap support (<50%) in this tree, the corresponding clade was supported by a bootstrap value of 98% in the former phylogenetic tree (Fig. 4A). TriNPC2.6 clade was composed only of proteins from *T. squamosa*.

**Fig. 4.**
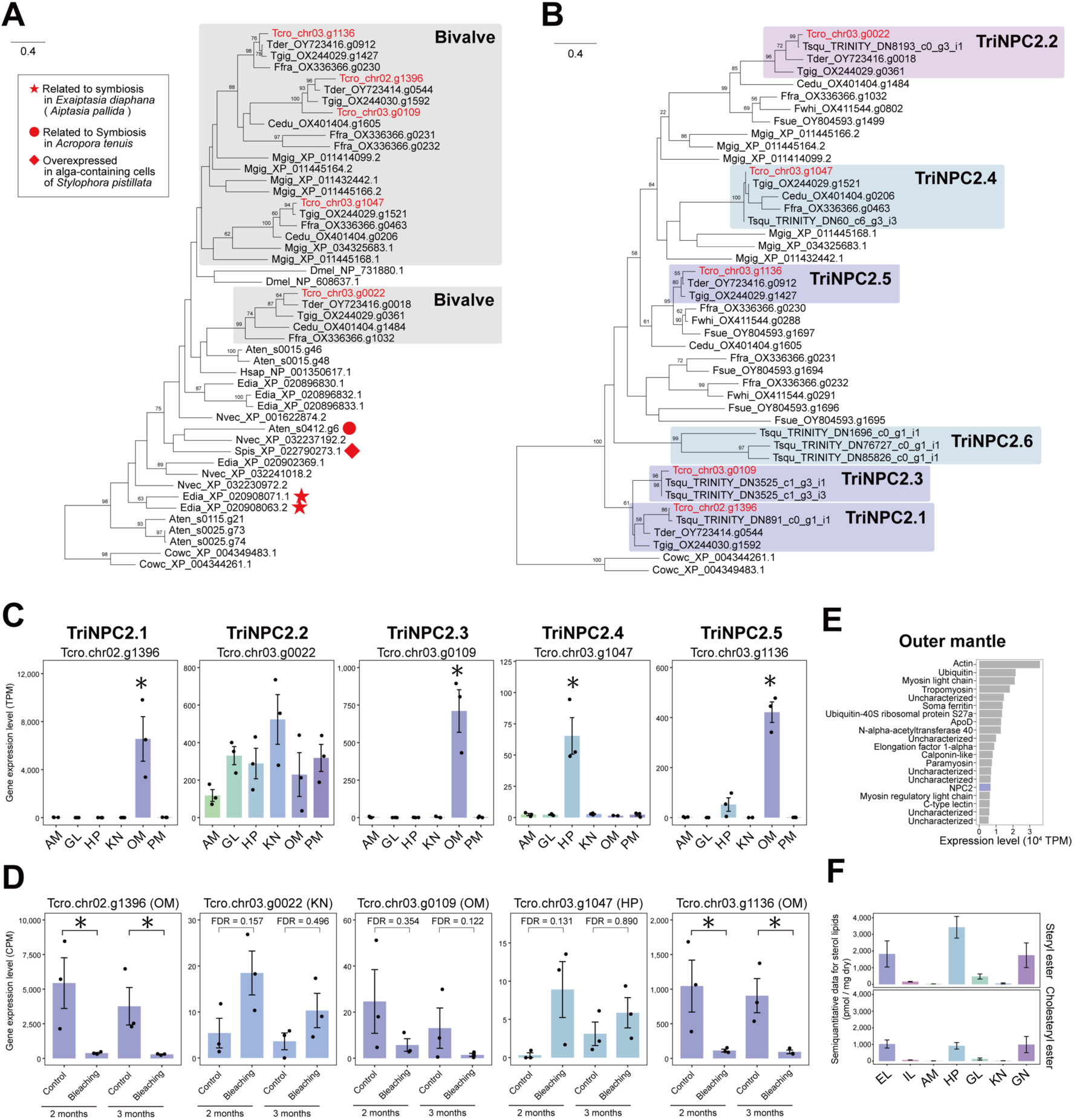
Phylogeny and expression patterns of genes related to sterol lipid transport: NPC2. (**A**) Molecular phylogenetic tree of NPC2 including cnidarians. A maximum likelihood molecular phylogenetic tree was constructed using 144 amino acid residues. Proteins from *T. crocea* are highlighted in red. Proteins suggested by previous studies to be involved in symbiosis with dinoflagellates in cnidarians (*50*, *54*, *56*, *57*) are highlighted by symbols. Bootstrap values > 50% are shown on branches. Taxonomy and protein IDs are separated by underscores. For proteins retrieved from NCBI GenBank, accession numbers are shown. (**B**) A molecular phylogenetic tree of NPC2 focusing on bivalves. The maximum likelihood molecular phylogenetic tree was constructed using 132 amino acid residues. Proteins from *T. crocea* are highlighted in red. Bootstrap values > 50% are shown on branches. For proteins retrieved from NCBI GenBank, accession numbers are shown. (**C**) Tissue-specific gene expression patterns of *T. crocea* NPC2 genes. (**D**) Gene expression responses of *T. crocea* NPC2 genes to dark-induced bleaching. (**E**) The 20 most highly expressed genes in outer mantle based on TPM. (**F**) Semiquantitative data of sterol lipids in *T. crocea* tissues. Lipid quantification data were obtained from Supplementary Table S1 of Sakai *et al.* (*59*), and values corresponding to sterol lipids (steryl esters and cholesteryl esters) were used for the construction of the bar plot. Steryl esters include campesterol, brassicasterol, β-sitosterol, and stigmasterol (phytosterols). Tcro, *T. crocea*; Tder, *T. derasa*; Tgig, *T. gigas*; Tsqu, *T. squamosa*; Ffra, *F. fragum*; Fwhi, *F. whitleyi*, Fsue, *F. sueziense*; Cedu, *C. edule*; Mgig, *M. gigas*; Dmel, *D. melanogaster*; Cowc, *C. owczarzaki*. AM, adductor muscle; GL, gill; HP, hepatopancreas (digestive diverticula); KN, Kidney; OM, outer mantle; PM, pedal mantle; GN, gonad; EL, epidermal layer of mantle (outer mantle); IL, inner tissue layer of mantle (inner mantle).

TriNPC2.1, TriNPC2.3, and TriNPC2.5 genes were highly expressed specifically in the outer mantle of *T. crocea*, whereas TriNPC2.4 was highly expressed in the hepatopancreas (Fig. 4C). In particular, TriNPC2.1 exhibited remarkably high expression levels in the outer mantle, with mean TPM values > 6,000, which ranks it among the 20 most highly expressed genes in this tissue (Fig. 4C and E). This value is comparable to those of several housekeeping genes, such as elongation factor 1-alpha and ribosomal proteins. It is likely that TriNPC2.1, 3, and 5 contribute to utilization of symbiont-derived sterols in the outer mantle, whereas TriNPC2.4 contributes to it in the digestive tract, where algal symbionts and filter-fed diets are digested. Indeed, large amounts of sterol lipids, including phytosterol lipids, were detected in the epidermal layer of the outer mantle, hepatopancreas, and gonad of *T. crocea* (Fig. 4F) (*59*).

TriNPC2.1 and TriNPC2.5 genes were significantly downregulated during bleaching, and TriNPC2.3 showed a similar trend, though the difference was not statistically significant (Fig. 4D). This may be a response to decreased sterol supply due to reduction of symbionts. On the other hand, TriNPC2.2 and TriNPC2.4 genes were not downregulated during bleaching, and their expression even appeared slightly elevated in the kidney and hepatopancreas, respectively, although not with statistical significance (Fig. 4D). This may reflect a shift in sterol acquisition and utilization pathways, including increased reliance on filter feeding and/or altered use of internal sterol lipids, compensating for reduced supply from symbionts. Overall, sterols are suggested to be a key component acquired from algal symbionts in the giant clam outer mantle. Sterol lipids are essential components for eukaryotic cells, but many marine invertebrates, including some molluscs and cnidarians, exhibit absent or weak de novo sterol biosynthesis and therefore rely on dietary sterols (*60*, *61*). Accordingly, sterol provisioning by algal symbionts represents a key nutritional benefit of algal-animal symbioses and is likely essential for giant clams, as well as cnidarians (*54*, *62*) and acoels (*63*).

### NRT2 Nitrate Transporter: An Ancient Plant-like Gene Co-opted for Photosymbiosis in Bivalves?

#### Evolutionary Origin of Animal NRT2

Among outer mantle-specific HEGs that were downregulated at 2 and 3 months during the bleaching experiment, we identified a DEG showing high sequence similarity to high-affinity nitrate transporter 2 (NRT2) of the model plant, *Arabidopsis thaliana* (Fig. 2D and 3E). However, a previous study concluded that NRT2 homologs are possessed only by autotrophic organisms such as plants and algae and in osmotrophs, e.g. fungi (*64*). Given this unexpected homology, we investigated the evolutionary origin of this gene (see Supplementary Text). The overall topology of the phylogenetic tree including known NRT2 homologs and candidate animal homologs was largely consistent with that reported by Ocaña-Pallarès *et al*. (*64*), with metazoan proteins, specifically from molluscs, annelids, and rotifers, forming a distinct clade within the eukaryotic NRT2 group, separate from bacterial homologs and supported by a bootstrap value of 100%. In the eukaryotic clade, a subgroup comprising animals, fungi, *Corallochytrium limacisporum* (Teretosporea, Opisthokonta), and *Labyrinthulea* was recovered with 76% bootstrap support (Fig. 5B). These results suggest conservation of NRT2 homologs in some animal lineages.

**Fig. 5.**
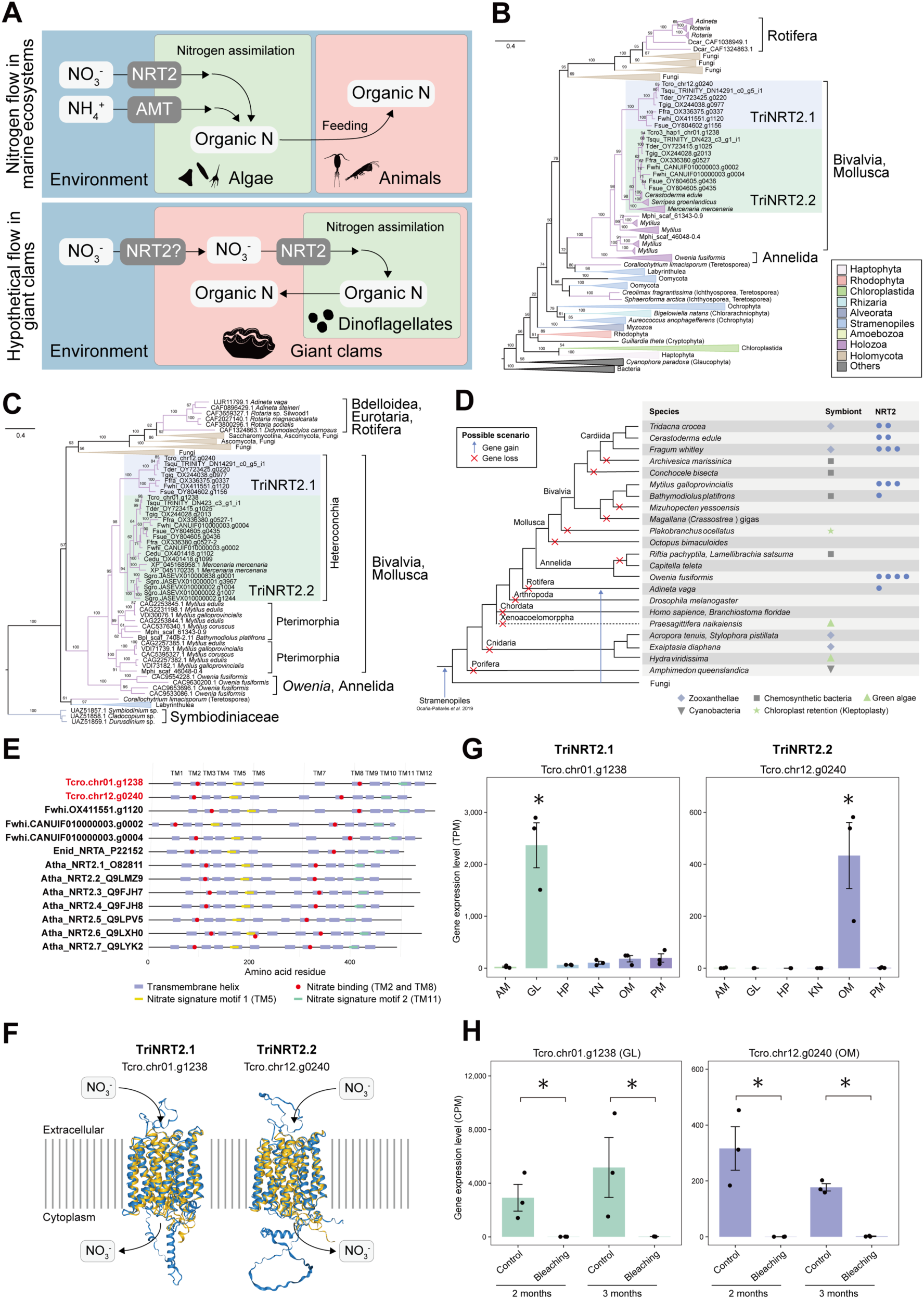
Phylogeny, domain structure, and gene expression pattern of NRT2 nitrate transporter, and the dark-induced bleaching experiment. (**A**) Schematic illustration of general nitrogen flow in marine ecosystems (top) and an example of hypothetical nitrogen flow in giant clams (bottom). (**B**) A molecular phylogenetic tree of animal NRT2 homologs with a broad eukaryotic NRT2 dataset adopted from Ocaña-Pallarès *et al*. (*64*). A maximum likelihood molecular phylogenetic tree was constructed using 430 amino acid residues. Bootstrap values > 50% are shown on branches. Taxonomic classification of eukaryotes is based on Ocaña-Pallarès *et al*. (*64*). (**C**) A molecular phylogenetic tree of animal NRT2 homologs with Symbiodiniaceae as an outgroup. Only homologs predicted to possess twelve transmembrane helices were retained as intact NRT2 homologs, and the longest transcript variant for each gene was used for the phylogenetic analysis. A maximum likelihood molecular phylogenetic tree was constructed using 418 amino acid residues. Bootstrap values > 50% are shown on branches. (**D**) Evolutionary history of animal NRT2. Examples of gene gain and loss timing, microbial symbionts (if applicable), and copy numbers of selected animals are shown. Horizontal gene transfer (HGT) from Stramenopiles to the common ancestor of Holozoa and Holomycota is the conclusion of Ocaña-Pallarès *et al*. (*64*). (**E**) Predicted domain structures of animal and plant NRT2. Transmembrane segments were predicted with DeepTMHMM 1.0. Residues labeled as “Nitrate binding” represent two arginine residues essential for substrate binding, according to Unkles *et al*. (*67*). The region labeled as the “Nitrate signature motif” corresponds to the conserved NS1 and NS2 motifs, along with surrounding, highly conserved regions identified by Unkles *et al*. (*68*), which are critical for nitrate transport. (**F**) Three-dimensional structure models of *T. crocea* NRT2 proteins (blue) predicted using the AlphaFold Server and reference proteins (*Chlamydomonas reinhardtii* NRT2.2 AF-Q39609-F1-model v4 for TriNRT2.1 and *Oryza sativa* NRT2.1 AF-P0DKG9 v4 for TriNRT2.2; yellow). Protein models are viewed from the plane of the plasma membrane. (**G**) Tissue-specific gene expression patterns of *T. crocea* NRT2 genes. *, tissue-specific HEGs. (**H**) Gene expression responses of *T. crocea* NRT2 genes to dark-induced bleaching. *, FDR < 0.05 (DEG). Tcro, *T. crocea*; Tder, *T. derasa*; Tgig, *T. gigas*; Tsqu, *T. squamosa*; Ffra, *F. fragum*; Fwhi, *F. whitleyi*, Fsue, *F. sueziense*; Cedu, *C. edule*; Sgro, *S. groenlandicus*; Mphi, *M. philippinarum*; Enid, *E. nidulans*; Atha, *A. thaliana*. AM, adductor muscle; GL, gill; HP, hepatopancreas (digestive diverticula); KN, Kidney; OM, outer mantle; PM, pedal mantle.

From the metazoan sequences obtained above, we selected only those predicted to contain 12 transmembrane helices. These sequences were considered putative intact NRT2 homologs, and were used for subsequent analysis. An additional search also identified a putative NRT2 homolog from a deep-sea mussel harboring chemosymbiotic bacteria, *Bathymodiolus platifrons*, while no clear homologs were found in other tested animals (Fig. 5D). Molecular phylogenetic analysis of these genes recovered a strongly supported clade comprising dinoflagellates (Symbiodiniaceae) and another clade of Opisthokonta + Labyrinthulea, both with 100% bootstrap support (Fig. 5C). In the latter, fungi + rotifers formed a clade that was sister to a mollusc + annelid clade, supported by bootstrap values of 94% and 100%, respectively. In the mollusc + annelid clade, bivalve sequences formed a monophyletic group with *Owenia fusiformis* (Annelida) proteins as their sister clade (100% support). The Heteroconchia-only clade, supported by a 98% bootstrap value, was further divided into two clades: Clade I (100% support) and Clade II (94% support). Notably, the former included sequences exclusively from *Tridacna* and *Fragum*, both of which host symbiotic dinoflagellates, while the latter included other heteroconchian bivalves in addition to the two genera. These results support the conservation of NRT2 homologs only in limited lineages of rotifers, bivalves, and annelids (Fig. 5C and 5D).

A previous study suggested that NRT2 homologs have undergone multiple horizontal gene transfer (HGT) events among eukaryotic lineages, and that one such event may have occurred from Labyrinthulea to the common ancestor of Opisthokonta (*64*). The present analysis supporting a single clade comprising animals (molluscs and annelids), Teretosporea, and Labyrinthulea (Fig. 5B) implies that the NRT2 homolog acquired at the ancestral stage of Opisthokonta has been retained in these animal lineages (Fig. 5D). In addition, repeated gene duplication and loss events appear to have occurred in the bivalve and annelid lineages, indicating a complex evolutionary history of this gene family, possibly associated with ecological diversification. Fig. 5D illustrates one of the possible evolutionary scenarios consistent with our results. This scenario assumes a minimal number of gene acquisition events and multiple gene losses, based on the assumption that loss of NRT2 genes could have occurred easily in animals, which do not assimilate nitrogen. Details of the search for animal NRT2 homologs and additional information are provided in the Supplementary Text and Fig. S7.

### Functional Implications of Giant Clam NRT2 in Relation to Photosymbiosis

All *Tridacna* species possess two copies of NRT2 genes, while *Fragum* species have been suggested to have three copies. Hereafter, the NRT2 of *Tridacna* in Clade I will be referred to as TriNRT2.1, and those in Clade II as TriNRT2.2. Copy number of NRT2 genes varied across animal species, from 0 to 5 (Fig. 5D). *T. crocea* TriNRT2.1 (Tcro.chr01.g1238) and TriNRT2.2 (Tcro.chr12.g0240) contained a conserved Major Facilitator Superfamily domain (Pfam: PF07690) and 12 transmembrane helices (TMs), consistent with the topology of plant NRT2 proteins (Fig. 5E). The three-dimensional structure of *T. crocea* NRT2 predicted by the AlphaFold Server (*65*) showed high structural similarity to algal and plant NRT2 proteins when compared using the Foldseek Server (*66*) (Fig. 5F and S9). Amino acid residues essential for the function of NRT2 homologs have been characterized in the fungus *Emericella* (*Aspergillus*) *nidulans*. Two arginine residues required for substrate binding (R87 in TM2 and R368 in TM8; Unkles *et al.* 2004) were completely conserved in *E. nidulans, T. crocea, F. whitleyi,* and *A. thaliana* (Fig. 5E and S8). In addition, other highly conserved and functionally important residues (*67*, *68*) were also identical in animal sequences to those of either *E. nidulans* or *A. thaliana* (Fig. S8). These conserved residues, together with overall structural similarity, strongly suggest that NRT2 homologs in giant clams possess nitrate transport activity comparable to that of fungal and plant NRT2 proteins.

*T. crocea* NRT2 homologs exhibit distinct expression patterns; TriNRT2.1 is a gill-specific HEG, while TriNRT2.2 is an outer mantle-specific HEG (Fig. 5G). Both were also significantly downregulated at 2 and 3 months during the dark-induced bleaching experiment (Fig. 5H), suggesting their possible involvement in symbiotic mechanisms.

Photosynthetic organisms assimilate nitrogen, unlike animals (Fig. 5A). Therefore, uptake of nitrate or ammonium from the environment is essential for plants and algae, and availability of inorganic nitrogen is one of the most important factors limiting their growth (*69*, *70*). NRT2 is a nitrate transporter responsible for nitrate uptake and transport (*71–73*). Previous studies that comprehensively surveyed NRT2 homologs among eukaryotic lineages concluded that such homologs are found only in autotrophic organisms, e.g., plants and algae, and in osmotrophs, e.g. fungi, whereas heterotrophic animals lack NRT2 genes (*64*). Considering that animals do not assimilate nitrogen, but instead obtain organic nitrogen compounds by consuming other organisms, this conclusion is entirely reasonable. However, our analysis revealed that a small subset of animals indeed possesses NRT2 homologs (Fig. 5D). Furthermore, transcriptomic data suggest that NRT2 homologs in *Tridacna* may contribute to the symbiotic relationship with dinoflagellates, as discussed above. Then, what might be their physiological function?

Because nitrogen is an essential nutrient for symbiotic dinoflagellates, the host must supply it (*2*). In corals, although multiple redundant pathways are thought to exist, a major route involves transfer of ammonium (NH_4_^+^) generated through host cell metabolism to symbionts (*2*, *74*). In contrast, the mechanism of nitrogen supply in giant clams remains unclear. NRT2 may accomplish this by mediating nitrate uptake from seawater and its transport within the host body (Fig. 6). Indeed, both giant clams and their symbionts acquire nitrate from the environment (*75*, *76*). Given that TriNRT2.1 is specifically expressed in the gill, it is likely responsible for nitrate uptake from seawater. Absorbed nitrate is presumably transported to the outer mantle via the hemolymph. On the other hand, TriNRT2.2 is specifically expressed in the outer mantle, suggesting that it may be directly involved in supplying nitrate to symbiotic dinoflagellates. Alternatively, this tissue may also take up nitrate directly from seawater.

**Fig. 6.**
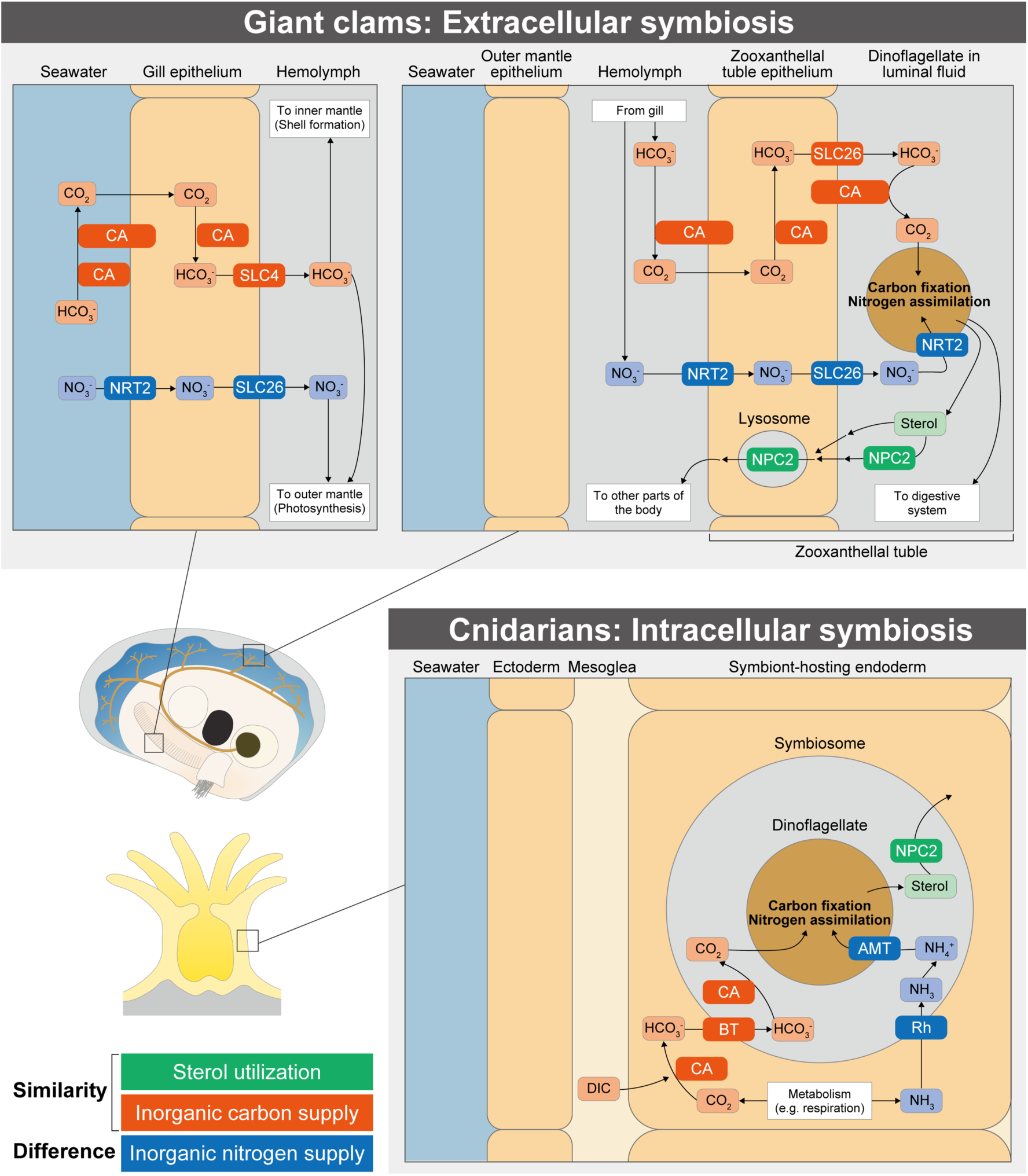
Schematic diagram of physiological functions of possible symbiosis-related proteins, focusing on CA, NPC2, and NRT2. These proteins were identified in this study and previous studies (*47*, *48*, *132*). Carbon and nitrogen supply and sterol utilization in corals proposed in previous studies are also shown (*54*, *74*, *133–135*). Acidification of the symbiosome in corals and the tubular lumen in giant clams, driven by V-type H^+^-ATPase (VHA), is thought to promote conversion of HCO_3_^−^ to CO_2_, but for simplicity this process is not depicted in the figure. In addition, nitrogen transport in the form of nitrate in corals and ammonium in giant clams is not depicted, but cannot be excluded. CA, carbonic anhydrase; DIC, dissolved inorganic carbon; BT, bicarbonate transporter; SLC, solute carrier protein; NPC2, NPC intracellular cholesterol transporter 2; NRT2, high affinity nitrate transporter 2; Rh, Rhesus channel; AMT, ammonia transporter.

NRT2 mediates transport of nitrate ions from the extracellular to the intracellular space (*71*, *72*). Symbiotic dinoflagellates reside extracellularly, in the zooxanthellal tubes of the outer mantle in giant clams. Therefore, to deliver nitrate to symbionts, a transporter capable of exporting nitrate from the host cytoplasm to the extracellular environment is required. A candidate for this function is a member of the SLC26A2-like protein family. Vertebrate SLC26A2 is known primarily as a sulfate transporter, but it is a multifunctional anion transporter that also exhibits nitrate efflux activity (*77*). The gene encoding this protein (Tcro.chr16.g1222) showed significantly decreased expression in the gill at 2 and 3 months, and also exhibited a downward trend in the outer mantle at 3 months (Fold change = 3.2, *p* = 0.0012, FDR = 0.089) (Fig. S10), suggesting its potential involvement in symbiosis. Based on these findings, we propose a model in which NRT2 on the apical membrane of gill epithelial cells acquires nitrate from seawater, SLC26A2-like releases it into the hemolymph, whereby it is then transported to the outer mantle. There, NRT2 in epithelial cells of zooxanthellal tubules imports nitrate into the cytoplasm, and SLC26A2-like subsequently exports it into the luminal fluid, allowing symbiotic dinoflagellates to absorb nitrate via their own NRT2 transporters (*78*) (Fig. 6).

If NH_4_^+^ is the primary form of inorganic nitrogen supplied to symbionts in corals, as NO_3_^−^ is in giant clams, this difference may reflect the contrasting nature of intracellular and extracellular symbioses. In corals, ammonia produced by host cellular metabolism is transported into symbiosomes. There, a low-pH environment, maintained by part of the CCM (V-type H^+^-ATPase), converts NH_3_ into NH_4_^+^ in a pH-dependent equilibrium. NH_4_^+^ can then be assimilated by symbionts (*74*) (Fig. 6). In giant clams, however, symbionts reside extracellularly in zooxanthellal tubules, requiring nitrogen to be transported into tubule lumens. Although a CCM is also expected in this compartment (*47*, *48*), it remains unclear whether nitrogen transport or pH regulation evolved first. If the former preceded the latter, early clam symbioses may have lacked an acidic microenvironment around symbionts, allowing highly toxic ammonia to accumulate. This would have posed a physiological risk to both the host and symbionts. Under these evolutionary constraints, nitrate, which is far less toxic (*79*, *80*), may have been favored as a safer inorganic nitrogen source for extracellular photosymbiosis.

### Shared and disparate molecular strategies of extracellular and intracellular photosymbiosis

In this study, we discovered the molecular basis of extracellular photosymbiosis in the boring giant clam, *Tridacna crocea*, by integrating chromosome-scale genome assembly and transcriptome analyses. Based on these results and previous findings, we propose a model illustrating the flow of key nutrients and metabolites between the host and symbionts in the clam body (Fig. 6). Our analyses reveal that NPC2 genes are highly expressed in the outer mantle, consistent with patterns observed in intracellular symbiosis of corals and other cnidarians. This indicates that a convergent molecular mechanism has been independently acquired in different symbiotic modes, suggesting that acquisition of symbiont-derived sterols provides a shared benefit to hosts in animal-dinoflagellate symbioses. In addition, expression of genes related to carbon concentrating mechanisms (CCM) support previous physiological and molecular studies, reinforcing the view that giant clams enhance photosynthesis by supplying inorganic carbon to their symbionts, like corals. In contrast, we found that NRT2 homologs, previously thought to be absent in animals, are present in the giant clam genome and likely mediate nitrate uptake and delivery to symbionts. This finding indicates that giant clams have a “plant-like” molecular trait that underlies their unique nutritional strategy, highlighting a key difference from corals in nitrogen transport. Expression of these genes decreased during bleaching, which suggests that the collapse of symbiosis triggers downregulation of nutrient transport pathways, underscoring their functional importance in symbiosis maintenance. Together, our findings imply mutual nutrient transport between the host and symbionts: sterol import from symbionts to the host and inorganic carbon and nitrogen supply from the host to symbionts. By contrasting shared and unique features between giant clams and corals, this study highlights both conserved and divergent strategies of photosymbiosis in marine invertebrates. These insights advance our understanding of evolution and physiology of animal–alga symbioses and provide a molecular and genomic basis for conservation of coral reef ecosystems, biodiversity maintenance, and sustainable management and aquaculture of giant clams.

## Materials and Methods

### Sample Collection for Genome Assembly and Gene Prediction

*Tridacna crocea* specimens for genome assembly and gene prediction were collected in Urasoko Bay, Ishigaki Island, Okinawa, Japan, under permits from Okinawa Prefecture (Numbers: 31-73, 2-60, and 3-66). Initially, collected clams were kept in an outdoor tank with a constant inflow of sand-filtered seawater at Yaeyama Station, Fisheries Technology Institute, on Ishigaki Island (Okinawa, Japan). A clam (individual ISG01) was then dissected with sterile scalpels. The gonad sample was kept at −80°C until use for genome sequencing.

For RNA-seq for gene prediction, eggs were collected from another adult clam (individual ISG03) and fertilized with sperm collected from two adult clams (individuals ISG04 and ISG05) following the method of (*59*, *81*). Fertilized eggs were kept in a 27°C incubator with 0.2-µm filtered seawater, and the water was replaced once a day. Larvae were collected 3, 24, and 72 h after fertilization. Eggs from individual ISG03 and larvae were quickly frozen in liquid nitrogen and kept at −80°C until use. Another adult clam (individual ISG02) was dissected, and adductor muscle, gill, gonad, kidney, hepatopancreas, and outer mantle were collected. Tissue samples were frozen and stored as described above.

Fertilized eggs were obtained using eggs from individual ISG06 and sperm from individuals ISG07 and ISG08. Following the method of (*82*), metaphase chromosomes from embryos were stained in VECTASHIELD Mounting Medium with DAPI (Vector Laboratories, USA) and observed under a ZEISS Axio Imager Z1 microscope (ZEISS, Germany).

### Genome and Transcriptome Sequencing

High-molecular-weight DNA was extracted from the sperm sample using a Nanobind Tissue Big DNA Kit (Circulomics, USA), and DNA size was assessed with a Femto Pulse system (Agilent Technologies, USA). Extracted DNA was sheared with Megaruptor 3 (Diagenode, USA), and then purified using AMPure PB (Pacific Biosciences, USA). Library preparation was conducted using a SMRTbell Express Template Prep Kit 2.0 (Pacific Biosciences). Size selection was carried out with BluePippin with High Pass Plus Gel Cassettes (Sage Science, USA) to remove library fragments smaller than 17 kb. HiFi sequencing was performed on a Sequel II system (Pacific Biosciences). The sequencing polymerase was bound to the SMRTbell library using a Sequel II Binding Kit 2.2, and the resulting library was loaded onto a SMRT Cell 8M and sequenced with a Sequel II Sequencing Kit 2.0, following the manufacturer’s protocol. A Hi-C library was prepared using a Dovetail Omni-C Kit (Cantata Bio, USA), following the manufacturer’s standard protocol. The resulting library was sequenced on a single lane of a NovaSeq 6000 system (Illumina, USA) using a NovaSeq 6000 SP Reagent Kit v1.5 (300 cycles; Illumina).

Total RNA was extracted using a POLYTRON Homogenizer (Kinematica, Switzerland) in combination with a Direct-zol RNA Miniprep Kit (Zymo Research, USA). RNA integrity was assessed using the High Sensitivity RNA ScreenTape assay for TapeStation systems (Agilent Technologies, USA). Intact poly(A)+ RNA was isolated with an NEBNext Poly(A) mRNA Magnetic Isolation Module (New England Biolabs, USA). An RNA-seq library was then prepared using an NEBNext Ultra™ II Directional RNA Library Prep Kit for Illumina and NEBNext Multiplex Oligos for Illumina (96 Unique Dual Index Primer Pairs) (New England Biolabs). 150-bp paired-end sequencing was then performed using a NovaSeq 6000 (Illumina, USA).

### Genome Assembly

Adaptor sequences in HiFi reads were filtered and trimmed with HiFiAdaptorFilt v.2.0.1 (*83*), and sequences longer than 20kb were used for the next step. The mitogenome was assembled with MitoHiFi v.3.0.0 (*84*) using the published mitogenome of *T. crocea* (NC_057530.1) (*85*) as a reference. For the nuclear genome, HiFi reads were assembled into contigs using Hifiasm v.0.19.7-r598 (*86*). We mapped HiFi reads to these contigs with minimap2 v.2.26-r1175 (*87*), and allelic contigs were identified and removed with Purge Haplotigs v1.1.2 (*88*). To identify sequences originating from the mitogenome and contaminants, we performed a homology search against the *T. crocea* mitochondrial genomic sequence (the one assembled in this study and OX031074.1) with BLASTN v.2.12.0+. In addition, we utilized FCS-GX v.0.4.0 (*89*) to identify sequences originating from contaminants. Contigs free of mitochondrial sequences and contaminants were scaffolded with YaHS v.1.2a.1 (*90*), incorporating Illumina shotgun sequences trimmed by TrimGalore v.0.6.10 (https://github.com/FelixKrueger/TrimGalore). Scaffolds were then manually curated using Juicebox v.2.15 (*91*).

Telomeric repeat sequences were identified using tidk v.0.2.41 with the option ‘--clade’ specified as ‘Cardiid’ (*92*). Completeness of the haplotype-phased draft genome was assessed with compleasm v.0.2.6 (*93*) using the metazoa_odb10 dataset. To assess the integrity of the draft genome, we conducted a BLASTN search against the mitochondrial genome and again performed a contaminant screening using FCS-GX. Additionally, a BLASTN search was performed against a custom database comprising *Symbiodinium microadriaticum* (GCA_001939145.1), *Symbiodinium* sp. (GCA_003297005.1), *S. kawagutii* (GCA_009767595.1), *S. natans* (GCA_905221605.1), *Cladocopium* sp. (GCA_003297045.1), *C. goreaui* (GCA_947184155.1), and *Durusdinium* sp. (*94*).

### Gene Prediction and Annotation

The assembled nuclear genome of *T. crocea* and genomes of *T. gigas*, *T. derasa*, *Fragum whitleyi*, *F. sueziense*, *Cerastoderma edule*, *Serripes groenlandicus*, downloaded from the NCBI GenBank database, were used for gene prediction (Table S1). Repetitive elements in scaffolds were identified *de novo* using a combination of RepeatScout v.1.0.6 (*95*) and RepeatMasker v.4.1.0 (http://www.repeatmasker.org). Repetitive elements were soft- and hard-masked based on length (>50 bp) and occurrence (more than 10 times). Gene prediction was first conducted with the BRAKER pipeline v.2.1.2 (*96*), with AUGUSTUS v.3.3.3, as in (*97*). RNA-seq reads generated in this study or downloaded from the NCBI SRA database (Table S5) were trimmed using TrimGalore v.0.6.10 (https://github.com/FelixKrueger/TrimGalore) with default settings, and aligned to the hard-masked genome sequence with HISAT v.2.1.0 (*98*). Then, alignment information was used for BRAKER gene prediction with options ‘UTR = on’, ‘softmasking’, and ‘AUGUSTUS_ab_initio’ with soft-masked genome sequence as input. We further conducted genome-guided transcriptome assembly to improve gene prediction using StringTie (*99*) with an option ‘-m 500’. Coding regions were then identified with TransDecoder v.5.7.1 (Haas, BJ. https://github.com/TransDecoder/TransDecoder). Gene models predicted by the above two strategies were compared with gffcompare v.0.12.6 (*100*). Genes that were absent in the AUGUSTUS prediction, but present in predictions from genome-guided transcriptome assembly or AUGUSTUS ab initio were added to gene predictions from AUGUSTUS using Another GFF Analysis Toolkit (AGAT) v.1.2.0 (Dainat J.https://www.doi.org/10.5281/zenodo.3552717). We named gene models using the pattern, [assembly ID]_chr[pseudochromosome number].g[number of genes in the pseudochromosome], or [assembly ID]_scf[scaffold number].g[number of genes in the scaffold]. The longest transcript variants of each gene were selected and translated into protein sequences with TransDecoder v.5.7.1. We assessed completeness of the proteome with BUSCO v.5.4.7 using the metazoan_odb10 dataset (954 genes) (*101*). In addition to *T. crocea*, gene predictions from other cardiid bivalve genome assemblies were also performed, as their gene models were not available from public databases.

The proteome was annotated with a BLASTP search (E-value cut off: 10^−5^) against the UniProt/SwissProt database (*102*) locally maintained on the NIG Supercomputer system (accessed on 2024-03-15) (Table S2). Domain structures were analyzed with InterProScan v.5.56-89 (*103*). In addition, eggNOG-mapper v2.1.12 (*104*) with eggNOG DB v5.0.2 was used as a complementary approach. For proteins of interest, transmembrane segments were predicted with DeepTMHMM 1.0 (*105*), and 3D structures were predicted using AlphaFold Server (*65*) and compared protein structure collections using the Foldseek Server (*66*).

Repetitive elements identified above were annotated as in (*42*). Briefly, sequences were searched against RepeatMasker.lib and RepeatPeps.lib bundled with RepeatMasker v.4.1.0 by BLASTN and BLASTX (E-value cut off: 10^−5^). The composition of repetitive elements was summarized with a perl script from RepeatMasker package (calcDivergenceFromAlign.pl), adding the class ‘Novel’ for non-annotated or possible novel repeat elements.

### Clustering of Orthologous Genes

In addition to *T. crocea* gene models predicted from the principal pseudohaplotype of the diploid genome assembled in this study, we used 21 publicly available metazoan gene models, including those of five cardiid bivalves symbiotic with Symbiodiniaceae algae, two non-symbiotic cardiid bivalves, two cnidarians symbiotic with Symbiodiniaceae algae, and a non-symbiotic cnidarian (Table S3). The longest transcript variants of each gene were selected and used to cluster orthologous groups (OGs) with OrthoFinder v.2.5.4 (*106*). OGs were considered gene families in this study.

### Molecular Phylogenetic Analysis and Divergence Time Estimation

For phylogenetic analysis of each gene, amino acid sequences or nucleotide sequences were aligned using MAFFT v.7.310 with the ‘-auto’ option (*107*), and gaps were removed using TrimAL v.1.4 with the ‘-gappyout’ option (*108*). Maximum likelihood analysis of aligned sequences was performed using RAxML v.8.2.12 with 100 bootstrap replicates and the ‘PROTOGAMMAAUTO’ option (*109*). Classification of species shown in phylogenetic trees follows the World Register of Marine Species (*110*).

For concatenated phylogenetic analysis, we used single-copy gene families. All amino acid sequences belonging to the same gene family were aligned using MAFFT v.7.310 with the ‘-auto’ option, and all gaps were removed using TrimAL v.1.4 with the ‘-nogaps’ option. All sequences from the same species were concatenated, and a maximum likelihood phylogenetic analysis was executed using RAxML v.8.2.12 with 100 bootstrap replicates and the ‘PROTOGAMMAAUTO’ option.

Divergence times of 22 metazoan species were estimated using the MCMCTREE module implemented in PAML v.4.8 (*111*), with the maximum likelihood phylogenetic tree constructed using concatenated amino acid sequences of 402 single-copy gene families as a reference tree. First, amino acid substitution rates were roughly estimated using the CODEML module implemented in PAML. Branch lengths, gradient vectors, and Hessian matrices were then calculated under the WAG+G substitution model. These were used for Bayesian divergence time estimation under the independent rates (IRs) relaxed clock model. Two priors, the overall substitution rate (rgene_gamma) and rate-drift parameter (sigma2_gamma), were set at G(1, 16.0) and G(1, 6.4). The burn-in, sample frequency, and number of samples were set to 1 million, 1,000, and 10,000, respectively. Six calibration points adopted from (*112*, *113*) were used (Table S4).

### Identification of NRT2 Gene Homologs

NRT2 gene homologs were searched with BLASTP against the 21 metazooan gene models and the NCBI non-redundant (nr) protein database locally maintained on the NIG Supercomputer system (accessed on 2024-11-07). *Tridacna crocea* putative NRT2 sequences (Tcro.chr01.g1238 and Tcro.chr12.g0240) were used as queries, and the *E*-value cutoff was set to 10^−5^. Animal proteins were extracted from BLASTP hits based on NCBI Taxonomy data, and molecular phylogenetic analysis was performed with proteins from the corresponding gene family (OG0007170) with datasets adopted from previous studies (*64*, *114*). Candidate animal NRT2 proteins were manually selected based on the topology of the phylogenetic tree, and used for further analyses. Putative NRT2 of *Bathymodiolus platifrons* (described below) and three genera of Symbiodiniaceae (*78*) were added. Additionally, proteins with fewer than 12 transmembrane regions were removed, and isoforms other than the longest one were also removed. Candidate proteins from *T. squamosa* were also added for both proteins. Molecular phylogenetic analysis was then performed as described above, but the mafft alignment strategy employed the “--localpair” option. Detailed information is provided in Supplementary Text and Fig. S7.

A BLASTP search was also performed against gene models of major animals that host photosynthetic or chemosynthetic symbionts; *Archivesica* (*Calyptogena*) *marissinica* (*115*) and *Conchocele bisecta* (*30*) downloaded from MolluscDB (*116*), *Bathymodiolus platifrons* (*117*), *Plakobranchus ocellatus* (GCA_019648995.1), *Riftia pachyptila* (*118*), *Lamellibrachia satsuma* (*119*), *Praesagittifera naikaiensis* (*120*), *Stylophora pistillata* (GCF_002571385.2), *Hydra viridissima* (*121*), and *Amphimedon queenslandica* (GCF_000090795.1). *Tridacna crocea* putative NRT2s were used as queries, and the *E*-value cutoff was set to 10^−5^.

### Tissue-specific Gene Expression Analysis of *T. crocea*

For tissue-specific transcriptome analysis, three clams (individuals M1-M3) were collected from Miyakojima City Center for Stock Enhancement, located on Miyako Island (Miyakojima, Okinawa, Japan). Immediately after collection, clams were washed with seawater filtered through a 0.22-µm Durapore hydrophilic PVDF membrane (Merck KGaA, Darmstadt, Germany). Specimens were dissected with disposable sterile scalpels, and tissue samples approximately 5 mm square were collected from six tissues: adductor muscle, gills, hepatopancreas, kidney, outer mantle, and pedal mantle (Fig. 2A). Tissue samples were preserved in RNAlater Stabilization Solution (Thermo Fisher Scientific, Waltham, Massachusetts, USA) and stored at −20°C until further use.

Total RNA was extracted from tissue samples using TRIzol reagent (Thermo Fisher Scientific, Waltham, Massachusetts, USA). Extracted RNA was cleaned with an RNeasy Mini Kit (QIAGEN) following the manufacturer’s instructions with DNase treatment. Quality of RNA was evaluated with a 4200 TapeStation and RNA ScreenTape (Agilent Technologies). RNA-seq was performed as described above.

RNA-seq reads obtained from three specimens were trimmed with TrimGalore v.0.6.10 (https://github.com/FelixKrueger/TrimGalore) and mapped to *T. crocea* gene models using STAR 2.7.11b (*122*). Expression levels were quantified using featureCounts, a part of Subread (*123*). Mapped counts were normalized by the trimmed mean of M values (TMM) method, and then converted to counts per million (CPM) using edgeR v.4.2.2 in R v.4.4.2. Before statistical testing, we removed genes with low expression levels using the filterByExpr function. Then TMM-normalized CPMs in each tissue were compared pairwise with other tissues to identify tissue-specific highly expressed genes (HEGs). Differential expression analysis was conducted using the quasi-likelihood (QL) pipeline in edgeR, applying generalized linear models (GLMs) and quasi-F tests for each pairwise comparison of tissues. Obtained *p*-values were adjusted using the Benjamini–Hochberg method in edgeR. When the gene expression level of the region was significantly higher (false discovery rate; FDR < 0.05 and log_2_ fold change > 1) than the other five tissues, genes were considered tissue-specific HEGs. Transcripts per million (TPM) were also calculated with edgeR, and used for PCA analysis and visualization as bar plots.

Functional enrichment analysis for UniProt keywords and Gene Ontology terms using tissue-specific HEGs was performed on the web platform *Database for Annotation, Visualization and Integrated Discovery* (DAVID) 2025 with default settings (*124*, *125*). UniProt IDs of BLASTP top hit for *T. crocea* gene models were used as background dataset, and those for tissue-specific HEGs were analysed. Gene models that had no homologies with sequences in the SwissProt database were excluded from the analysis.

### Dark-induced Bleaching Experiment and Gene Expression Analysis

Reduction of symbiotic algae of giant clams can be induced by long-term exposure to darkness (dark-induced bleaching) (*27*, *28*). To investigate the impact of dark-induced bleaching on the bacterial community of *T. crocea*, a dark exposure experiment was conducted using the method of Uchida *et al.* (*28*). We utilized residual tissue samples to examine the host’s transcriptomic response by RNA-seq. Briefly, clams collected in Urasoko Bay were divided into two outdoor tanks with a constant inflow of sand-filtered seawater. Each tank was assigned to one of the two experimental groups: light or dark conditions. The former group was exposed to natural sunlight, while the latter was kept in continuously shaded conditions. Sampling was conducted twice, after approximately two and three months. Three individuals per condition were collected (biological replicates, n=3). Clams were dissected immediately using a scalpel. Tissue pieces were quickly frozen in liquid nitrogen, and stored at −80°C until use. The detailed experimental setup is explained in Uchida *et al*. (*28*).

Total RNA was extracted with TRIzol reagent (Thermo Fisher Scientific) and cleaned using an RNeasy Mini Kit (QIAGEN) with DNase treatment. A Collibri 3′ mRNA Library Prep Kit for Illumina (Thermo Fisher Scientific) was used for sequencing library preparation. Sequencing adaptors were attached by PCR amplification with 16 cycles of annealing, according to the manufacturer’s protocol. Each library was sequenced on a NovaSeq X Plus (Illumina) with 150-bp, paired-end reads. Reads 2 were discarded, and reads 1 were trimmed with TrimGalore, mapped with STAR, quantified with featureCounts, and statistically analyzed with EdgeR. Since 3′ mRNA-seq was performed, TPM was not calculated. Differentially expressed genes (DEGs) were identified by comparisons between the same tissue from the bleached group and control group at the same time points. Genes with considerable expression levels in each tissue and time point were extracted using the filterByExpr function in edgeR, and these gene sets were used as background datasets for functional enrichment analysis with DAVID.

### Tridacna squamosa Gene Prediction

For comparison with genome-sequenced tridacnid clams, protein-coding genes of *T. squamosa* were predicted using publicly available RNA-seq data. RNA-seq reads were trimmed with TrimGalore v.0.6.10 (https://github.com/FelixKrueger/TrimGalore). 10% each of seven samples (Table S18) were subsampled with SeqKit v.2.6.0 (*126*) and *de novo* assembled using Trinity v.2.15.1 with minimum contig length set to 200 bp (*127*, *128*). Assembled contig sequences were clustered using CD-HIT-EST v.4.8.1 with 90% sequence identity (*129*, *130*). Possible contaminant sequences were removed and remaining transcripts were translated into amino acid sequences in the same way as described above. The proteome was annotated with BLASTP searches against the UniProt/SwissProt database and InterProScan as described above. Only genes with functional annotation (20,680 gene models) were used for subsequent analysis.

## Acknowledgements

We thank Dr. Steven D. Aird for editing the manuscript, and Dr. Yuki Yoshioka for insightful discussions. Computations were partially performed on the NIG supercomputer at ROIS National Institute of Genetics.

## Author contributions

Conceptualization: TU, HY, CS

Data curation: TU

Formal analysis: TU

Funding acquisition: TU, HY, ES, CS

Investigation: TU, ES, MK, HY

Methodology: TU, ES, HY

Resources: GS, HY

Supervision: CS

Visualization: TU

Writing—original draft: TU

Writing—review & editing: TU, HY, GS, MK, ES, CS

## Data and materials availability

All data are available in the main text, the supplementary materials, or public database. All PacBio and Illumina reads are available under BioProject accession numbers PRJDB35853, PRJDB35854, PRJDB37588, and PRJDB37594, with DRA Run accession numbers DRR752275-DRR752279, DRR751819-DRR751828, DRR751931-DRR751948, and DRR752505-DRR752575. Genome assemblies are available under BioProject accession IDs PRJDB35682 (haplotype 1) and PRJDB35683 (haplotype 2). FASTA and GTF files of gene models are available at UTokyo Repository (https://doi.org/10.15083/0002014090). Scripts used for computational analyses are available at the GitHub repository (https://github.com/uchida-taiga/Tridacna-crocea_genome).

## Supplementary Materials

Supplementary materials include:

Supplementary Texts

Figs. S1 to S10

Legends for Tables S1 to S18

References

