## Supplementary Materials for "Genomic insights into photosymbiosis in giant clams: Comparisons with coral strategies"

Taiga Uchida *et al.*

**This PDF file includes:**

Supplementary Texts
Figs. S1 to S10
Legends for Tables S1 to S18
References

### Supplementary Texts

#### Apolipoprotein D (ApoD)

Outer mantle-specific highly expressed genes (HEGs) include four copies of ApoD genes. Two of them showed remarkably high expression levels, with mean TPM values > 2,900 and 17,000 (Fig. 4E and S4). Notably, as revealed by single-cell RNA sequencing, overexpression of ApoD genes also characterizes alga-hosting cells of stony corals, in concert with overexpression of NPC2 genes (*1*, *2*). In giant clams, one copy of ApoD was downregulated at the 3-month time point during bleaching, while another was upregulated at 2 and 3 months (Fig. 2B and 3D). These findings imply that ApoD is another key symbiosis-related gene common to corals and giant clams.

ApoD is a lipocalin-type apolipoprotein associated with HDL, but despite its various functions, it shows weak or undetectable binding to free cholesterol (*3*). Thus, it is unlikely to serve as a primary carrier of phytosterols derived from algal symbionts. Instead, ApoD may mediate short-range transport and quality control of other algal-derived lipids or their metabolites by solubilizing them and quenching lipid hydroperoxides. Because photosynthesis elevates the $O_2$ concentration in zooxanthellal tubules, the outer mantle epithelium likely faces high oxidative stress (*4*); ApoD's antioxidant activity may therefore be important for maintaining membrane homeostasis at this interface. Different responses of ApoD gene copies to bleaching may reflect functional diversification among paralogs.

#### Carbonic anhydrase (CA)

Like cnidarians, giant clams are thought to possess a carbon-concentrating mechanism (CCM) that enhances the supply of inorganic carbon ($CO_2$) to symbiotic dinoflagellates (*4*–*6*). Among key components of the CCM, several carbonic anhydrases (CAs) have been identified in *Tridacna squamosa*: a dual-domain CA (DDCA) localized at the host apical membrane of ctenidial (gill) epithelial cells (*7*), a cytosolic CA in tubule epithelial cells of the outer mantle (*8*), and a plasma-membrane-associated CA in symbiotic dinoflagellates (*9*). In the case of DDCA (GenPept accession ATC20500.1), a previous study predicted two transmembrane regions and a C-terminal GPI-anchor site. However, when the sequence was re-evaluated using DeepTMHMM 1.0.44 and DeepLoc 2.1, only a GPI-anchor signal was detected, suggesting that both CA domains likely face the extracellular side.

In the present study, twelve genes encoding CAs were identified in the *T. crocea* genome, all of which contain a Eukaryotic-type carbonic anhydrase domain (PF00194). Among these, Tcro.chr10.g1048 possesses one transmembrane helix (amino acid positions 1026–1035) according to DeepTMHMM, and its CA domain (positions 1052–1288) is predicted to be extracellular. Tcro.chr16.g0237, which corresponds to the DDCA, is predicted to be GPI-anchored.

Fig. S5 shows subcellular classification (cytosolic or extracellular) based on the CA domain position and tissue-specific expression profiles. Among carbonic anhydrases, three genes with particularly high expression levels (TPM > 500) are considered strong candidates for involvement in the CCM (shown below). Incorporating these findings with those of Mani *et al.* (*9*), we revised the CCM model proposed by Ip & Chew (*4*) as illustrated in Fig. 6.

- Tcro.chr10.g1048: Transmembrane protein with its CA domain oriented toward the cytoplasmic side; highly expressed in both gill and outer mantle.
- Tcro.chr11.g1134: Cytosolic protein; an outer mantle-specific HEG, but also showing high expression in the gill.
- Tcro.chr16.g0237: GPI-anchored protein with two extracellular CA domains; highly expressed in the gill (HEG).

**Identification of animal NRT2 homologs**

We searched for NRT2 homologs using three steps.

**Step 1**

- Protein sequences of *T. crocea* putative NRT2 were used as queries in BLASTP searches (e-value < $1\times10^{-5}$) against the NCBI nr database maintained on the NIG supercomputer (accessed on November 7, 2024). Based on NCBI taxonomic information, only sequences derived from animals were extracted from these hits.
- Proteins included in the same orthogroup as *T. crocea*'s putative NRT2 were added to the dataset.
- Previously reported NRT2 homologs of eukaryotes and prokaryotes described in Ocaña-Pallarès *et al.* (*10*) were also included.

All sequences were aligned using MAFFT, trimmed with trimAl, and poorly aligned sequences were manually removed. A maximum-likelihood phylogenetic tree was then constructed using RAxML. The resulting tree is shown in Fig. S7.

**Step 2**

- Based on topology of the phylogenetic tree obtained in **Step 1** and annotation of each sequence in the NCBI database, sequences that appeared to be contaminants, derived from microorganisms, were removed:

  XP_022835667.1, XP_003087464.1, XP_023254538.1 — these entries have already been removed from the latest version of nr database because they were determined to be contaminants.

  CAD7643483.1 — this sequence was predicted from a very short contig (~16 kb) in a draft genome. A BLASTN search against the nr database using the contig as a query showed 100% query coverage, 97% identity, and an e-value of 0 against *Acinetobacter* sp., suggesting bacterial contamination.

  EDO28285.1 — previously reported as a contaminant by Ocaña-Pallarès *et al.* (*10*).

  ETN62273.1 — the branch length was extremely long, and a BLASTP search against SwissProt retrieved non-NRT2 animal proteins such as Q9V7S5.1 (putative inorganic phosphate co-transporter, *Drosophila melanogaster*) and Q9NRA2.2 (Sialin, *Homo sapiens*) as top hits.
- Other animal-derived sequences from rotifers, mollusks, and annelids were retained because their within-clade topologies roughly matched known species relationships, and several were predicted from chromosome-level genomes assembled using long-read sequencing, indicating that contamination was unlikely.
- A putative homolog from *T. squamosa* was then added to the dataset.

A refined molecular phylogenetic analysis was then conducted (Fig. 5B).

**Step 3**

Based on the results of **Steps 1 and 2**, we concluded that certain animals possess NRT2 homologs. To investigate their taxonomic distribution in detail, we performed an additional analysis as follows:

- From the **Step 2** dataset, we extracted sequences from animals, fungi, *Corallochytrium limacisporum* (Teretosporea), and Labyrinthulea.
- For animal sequences, only the longest variant from each gene was retained, in order to avoid redundancy among transcript isoforms.
- A newly identified NRT2 homolog from the deep-sea mussel, *Bathymodiolus japonicus* (found via additional BLAST searches), was added.

NRT2 sequences from three clades of Symbiodiniaceae (*11*) were also included as an outgroup.

A final maximum-likelihood phylogenetic analysis was performed (Fig. 5C), and the resulting topology was used to construct the schematic summary shown in Fig. 5D.

**Evolutionary origin of rotifer NRT2 homologs**

In the phylogenetic analyses shown in Fig. 5B and 5C, NRT2 homologs from rotifers formed a distinct clade. Furthermore, this clade grouped with the Ascomycota, and that combined clade was nested within the fungal clade. All these clades were supported by high bootstrap values. From this topology, a horizontal gene transfer (HGT) event from fungi (likely Ascomycota) to rotifers is inferred. Additionally, because the animal NRT2 clade composed of mollusks and annelids does not include rotifer NRT2 homologs, it is plausible that rotifers initially lost the NRT2 gene and subsequently reacquired it via HGT. Indeed, rotifers exhibit an exceptionally high frequency of gene horizontal transfer; thus, a transfer event from fungi is not implausible (reviewed by Wilson *et al.* (*12*)).

**Protein structure of animal NRT2 homologs**

Compared with algal and plant NRT2s, the *T. crocea* NRT2 was predicted to have a longer and more flexible cytoplasmic domain between the sixth and seventh transmembrane helices. This feature is shared with fungal NRT2 homologs (NRTA or CRNA) (Fig. 5E, 5F, S9). Fungal NRT2 homologs, unlike those of plants, can transport nitrate independently of the accessory protein, NAR2, and this long cytoplasmic loop has been suggested to contribute to such NAR2-independent activity (*13–15*). Because no protein-coding gene showing homology to NAR2 was found in the *T. crocea* genome, it is likely that NRT2 in giant clams also functions independently, as in fungi.

**Other routes of inorganic nitrogen transport**

It is also possible that as in corals, giant clams possess multiple pathways to supply inorganic nitrogen. In bivalves, excretion of $NH_3$, a toxic byproduct of host metabolism, is mediated by Rh proteins (Thomsen et al., 2018). In corals, Rhesus channels (Rh proteins) localized on the symbiosome membrane transport $NH_3$ into the symbiosome lumen. The acidic environment inside symbiosomes, maintained as part of the CCM, shifts the chemical equilibrium toward $NH_4^+$, which can then be assimilated by symbiotic dinoflagellates (Thies et al., 2022). In giant clams, an Rh protein gene (Tcro.chr06.g1154) was significantly upregulated in the adductor muscle, gill, gonad, kidney, and outer mantle during bleaching (Table S2 and S13). This upregulation may reflect increased excretion of ammonia produced by catabolism of host proteins under nutrient-depleted conditions. Alternatively, it could indicate enhanced excretion of excess ammonia that has accumulated because symbionts no longer consume ammonium during bleaching. Another Rh protein gene, Tcro.chr01.g1185, showed high expression levels in the gill under normal conditions, but was also expressed at higher levels in the outer mantle compared to other tissues (Table S2 and S6). This suggests that in addition to excreting ammonia at the gill, this gene may also contribute to ammonium supply to symbiotic dinoflagellates, a mechanism similar to that in corals.

### Supplementary Figures

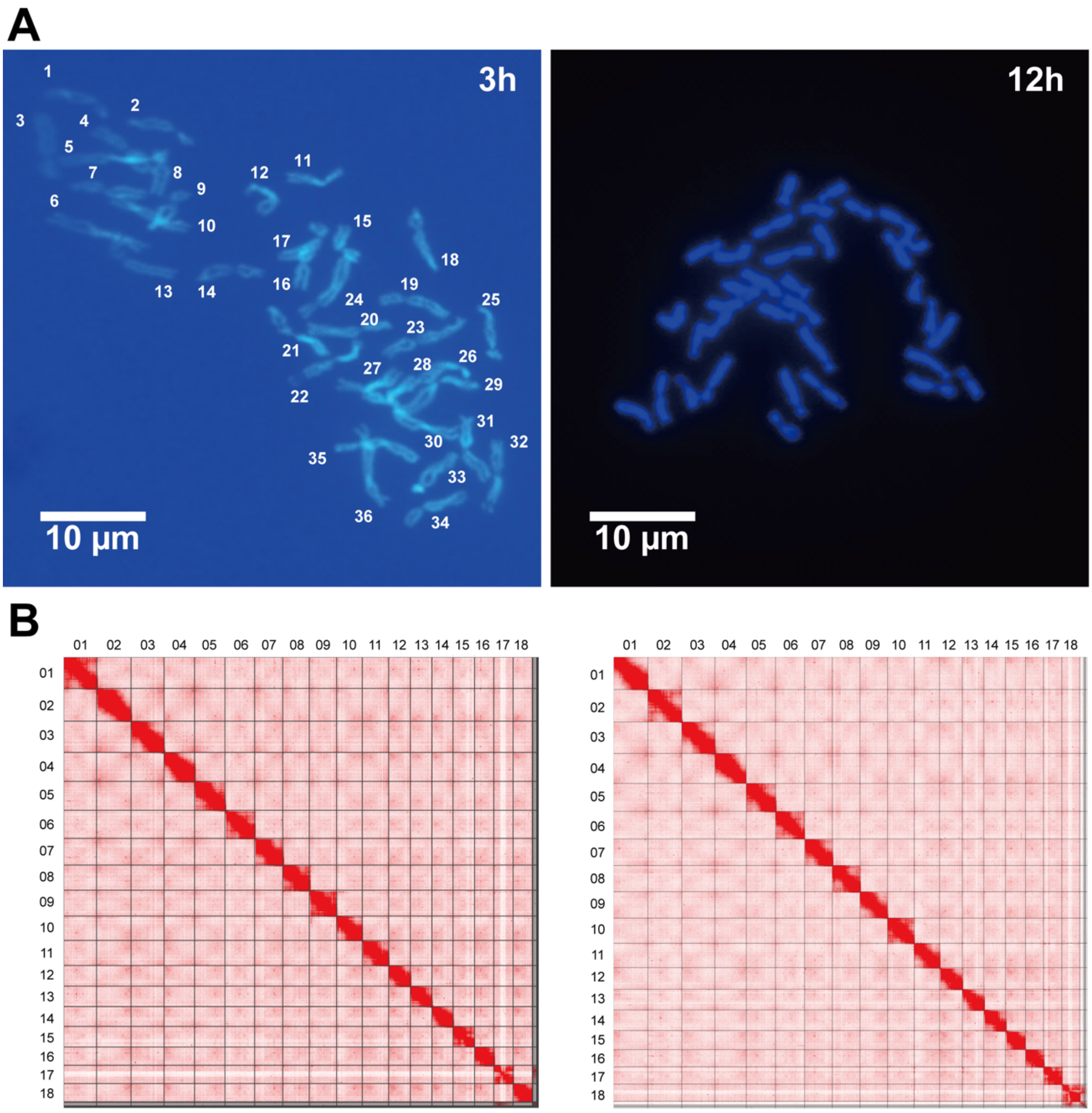


**Fig. S1.**

Additional information about genome assembly. (**A**) Microscopic observation of chromosomes from embryos of *T. crocea* 3 h (left) and 12 h (right) after fertilization. Numbers do not correspond to the assembled chromosome number. (**B**) Hi-C contact map of haplotype 1 (left) and 2 (right).

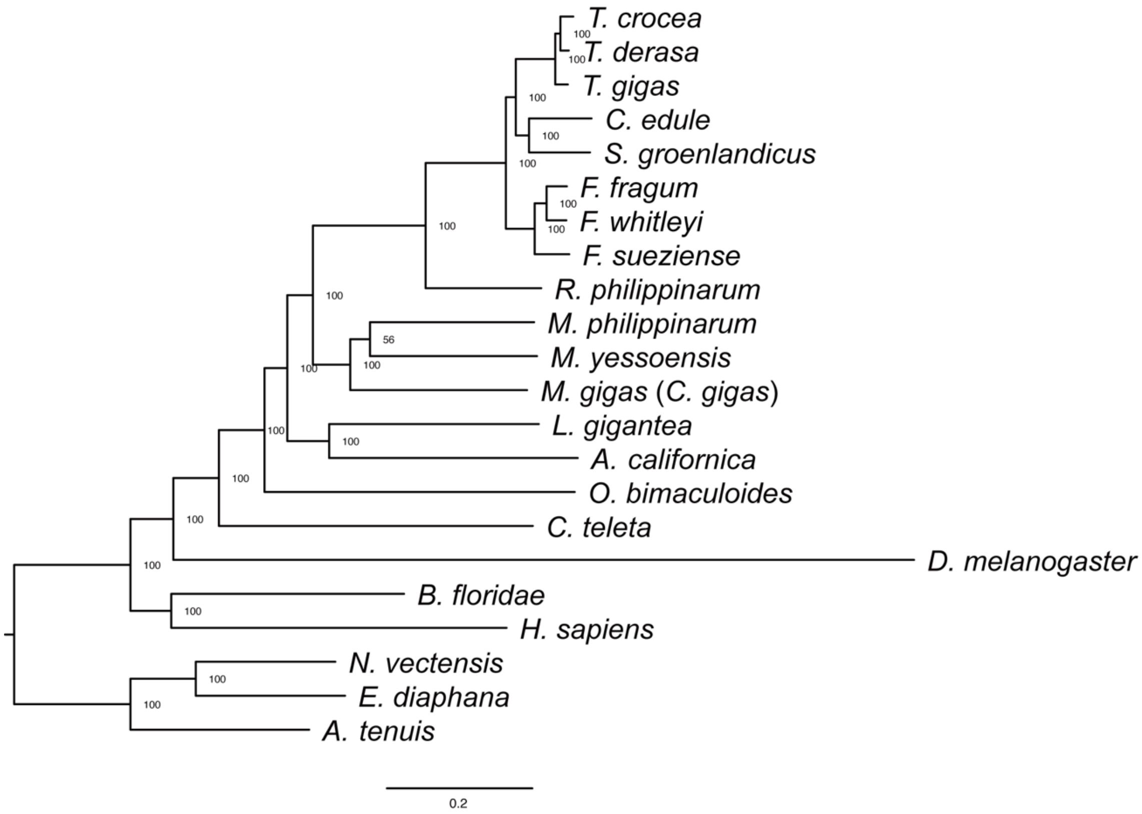


**Fig. S2.**

Maximum likelihood molecular phylogenetic tree of 22 metazoans with bootstrap values.

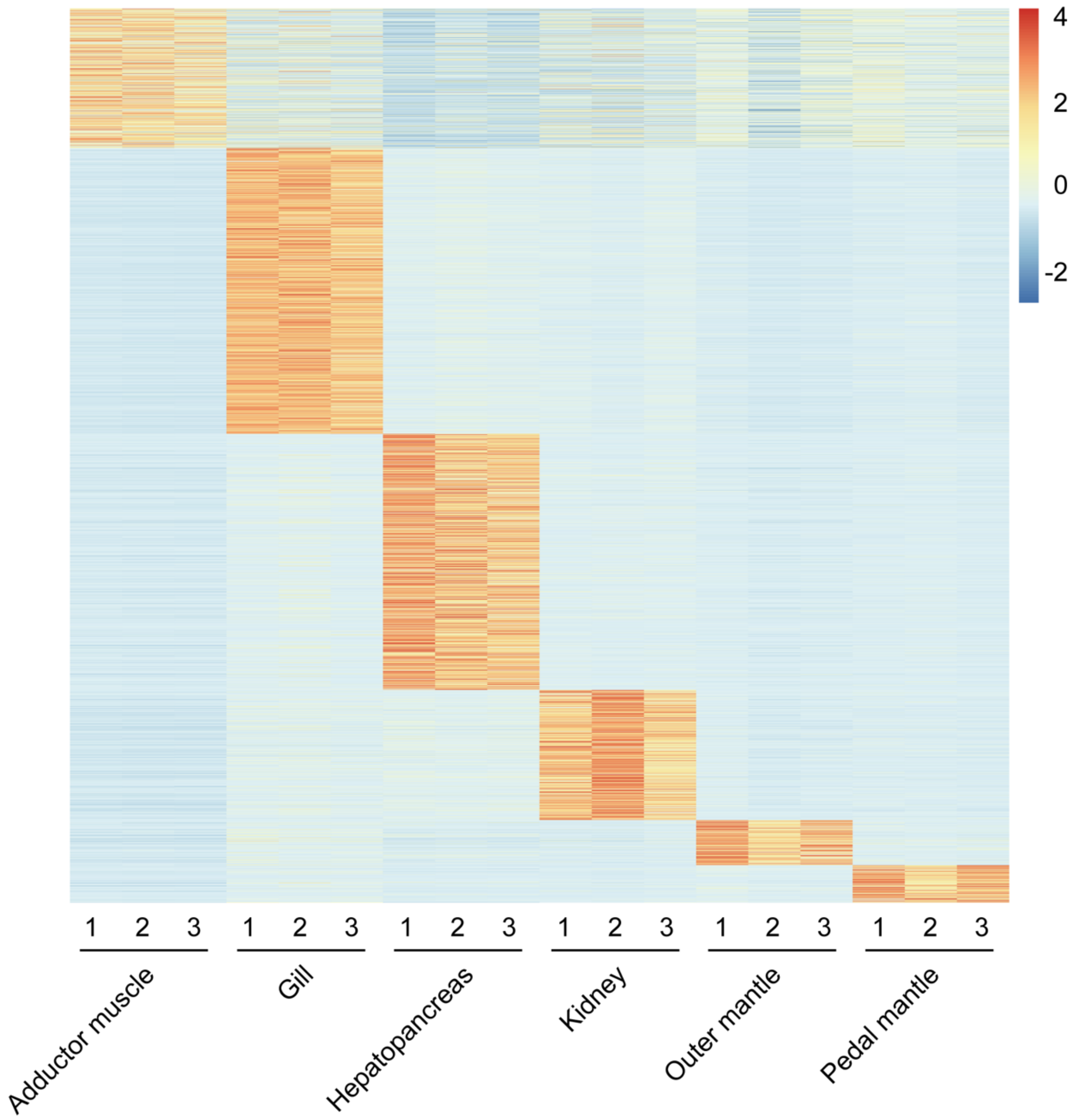


**Fig. S3.**

Heat map of relative expression levels (row Z-score transformed TPM) of tissue-specific DEGs. The TPM of each individual (n=3) is shown.

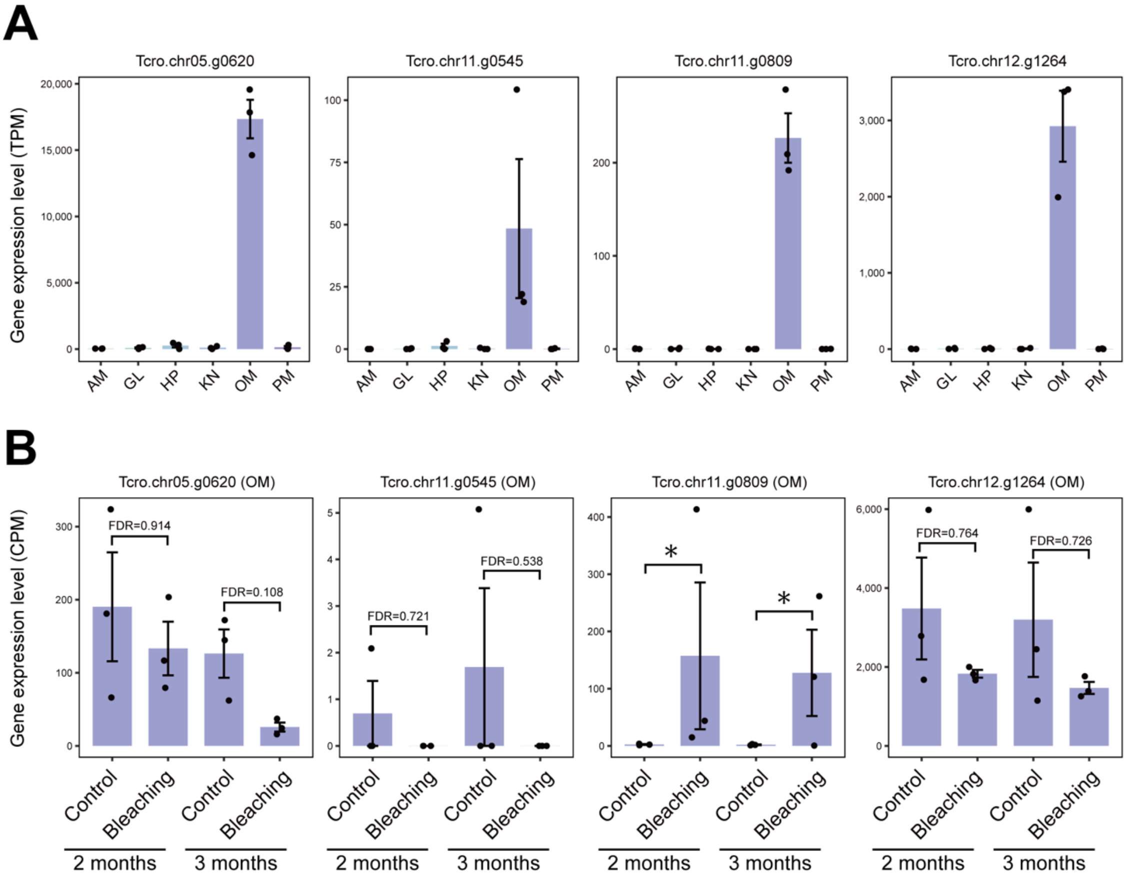


**Fig. S4.**

ApoD genes that were highly expressed in the outer mantle. (**A**) Tissue-specific gene expression. (**B**) Dark-induced bleaching experiment. * DEGs.

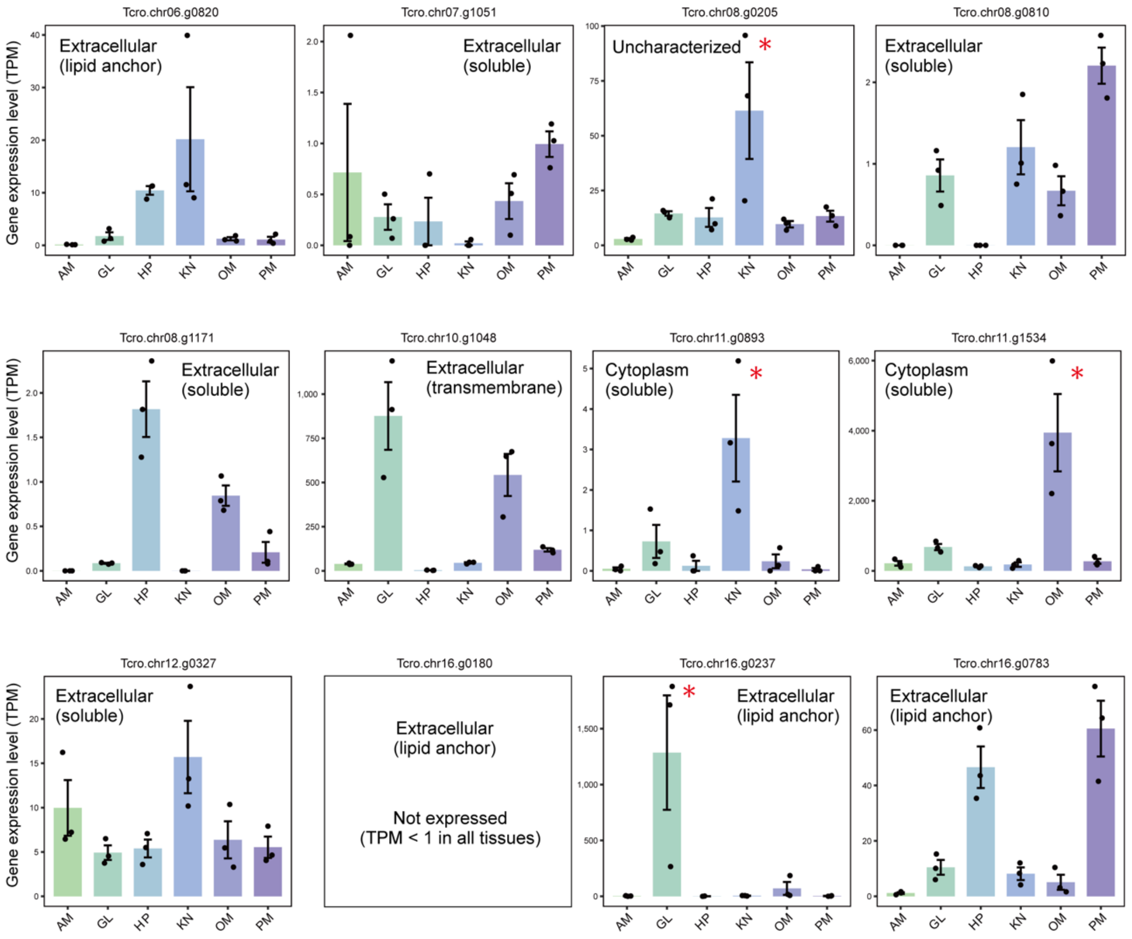


**Fig. S5.**

Gene expression, subcellular localization (soluble, lipid anchor, or transmembrane) and CA domain position (cytoplasm or extracellular) predicted by DeepTMHMM 1.0.44 and DeepLoc 2.1. * Tissue-specific HEGs or DEGs.

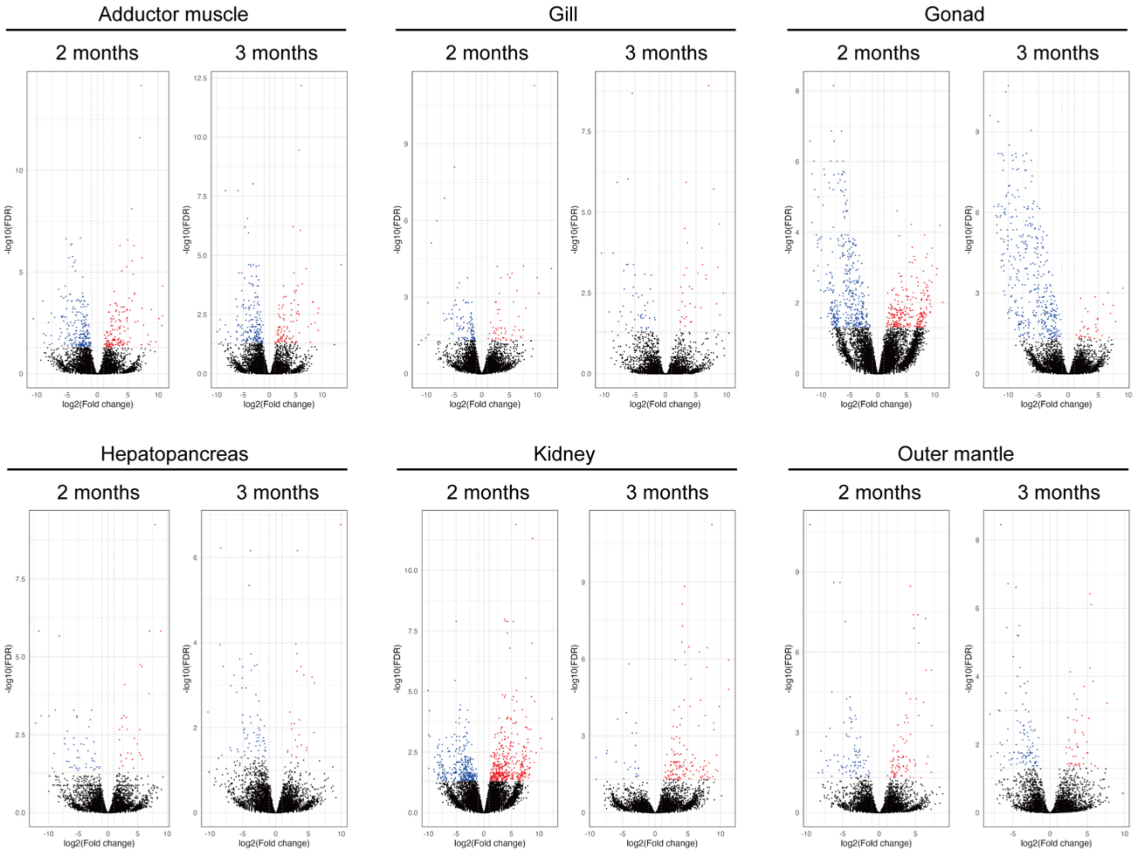


**Fig. S6.**

Volcano plot summarizing transcriptomic analysis after a dark-induced bleaching experiment. Upregulated DEGs are shown in red, and downregulated DEGs in blue.

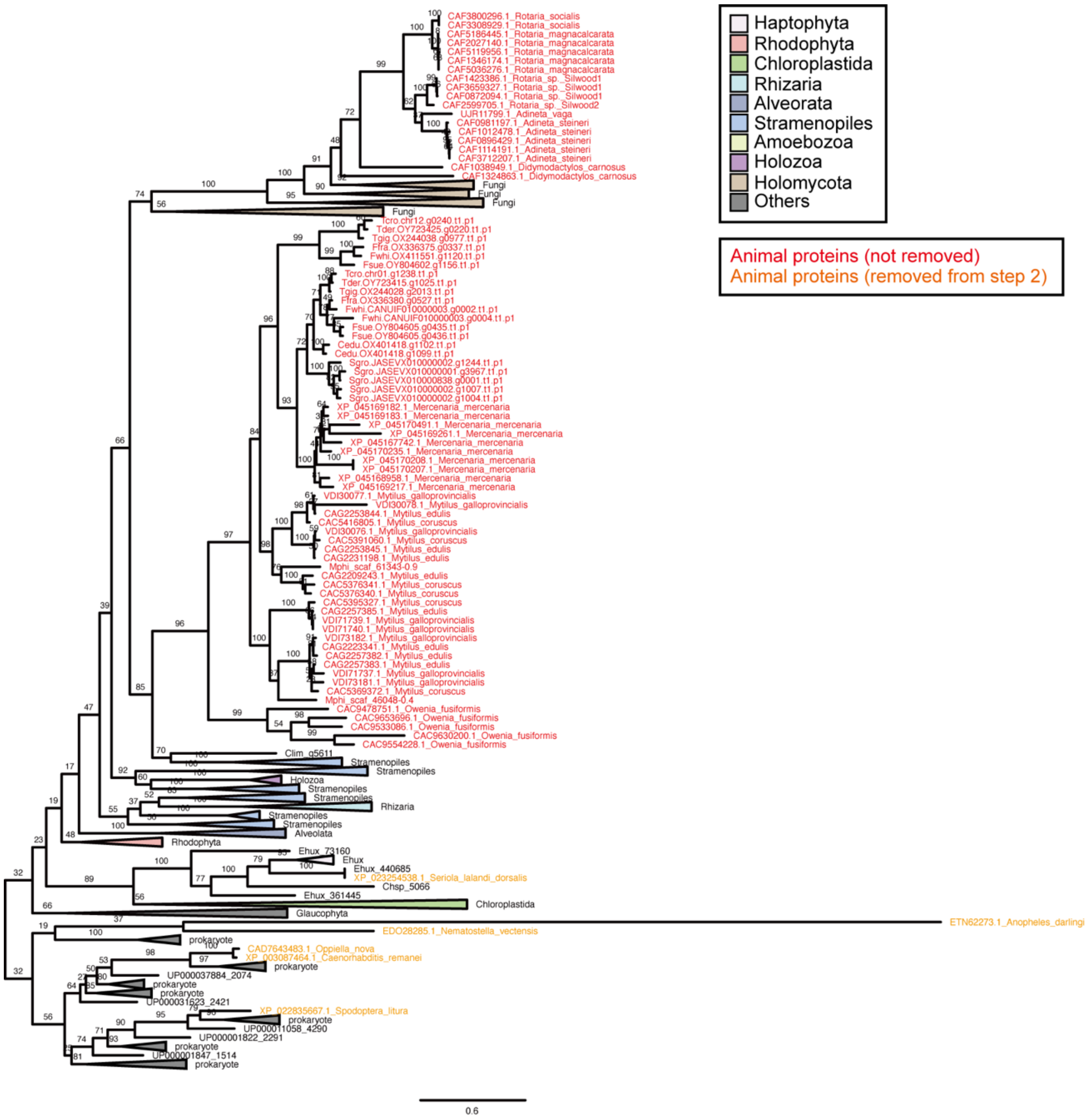


**Fig. S7.**

Maximum likelihood molecular phylogenetic tree of NRT2 homologs and candidate animal genes (the result of step 1 of the search). Numbers on the branches indicate bootstrap values.

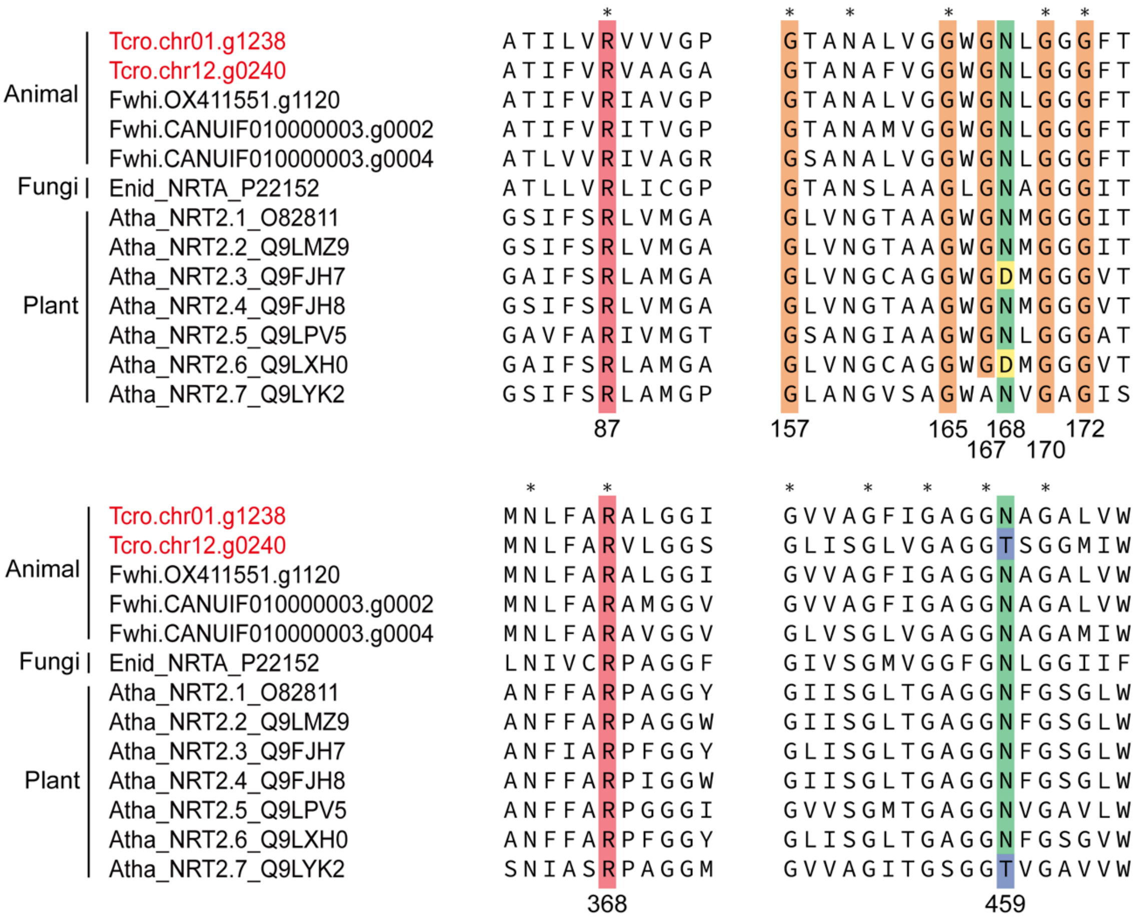


**Fig. S8.**

Alignment of amino acid sequences of NRT2 homologs. Residue number is based on *E. nidulans* NRTA (CRNA). Amino acid residues important for transporter activity (*16*, *17*) are highlighted in red, orange, or green.

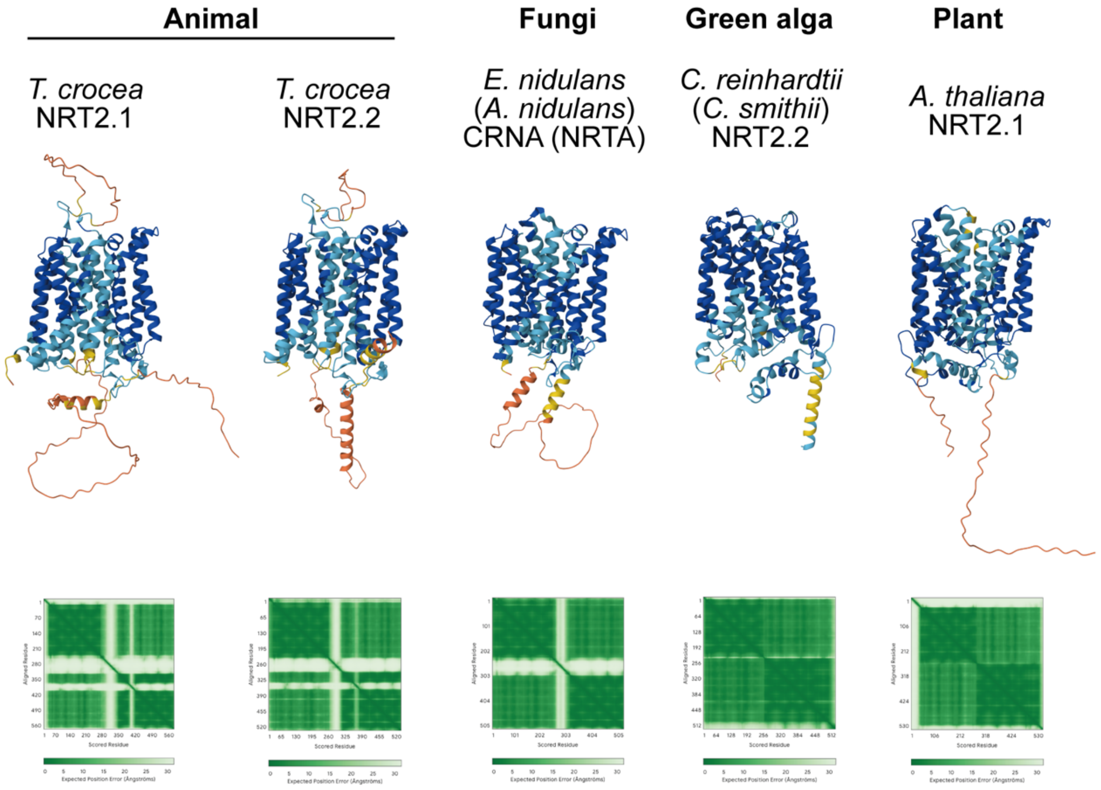


**Fig. S9.**

AlphaFold server outputs, in addition to Fig. 5F. Although the predicted transmembrane structures appear to differ between *T. crocea* + *C. reinhardtii* and *E. nidulans* + *A. thaliana*, this difference likely reflects whether the top-ranked model was oriented with the opening toward the extracellular or intracellular side. It should be noted that the orientation of the predicted structure cannot be specified in advance in the AlphaFold server.

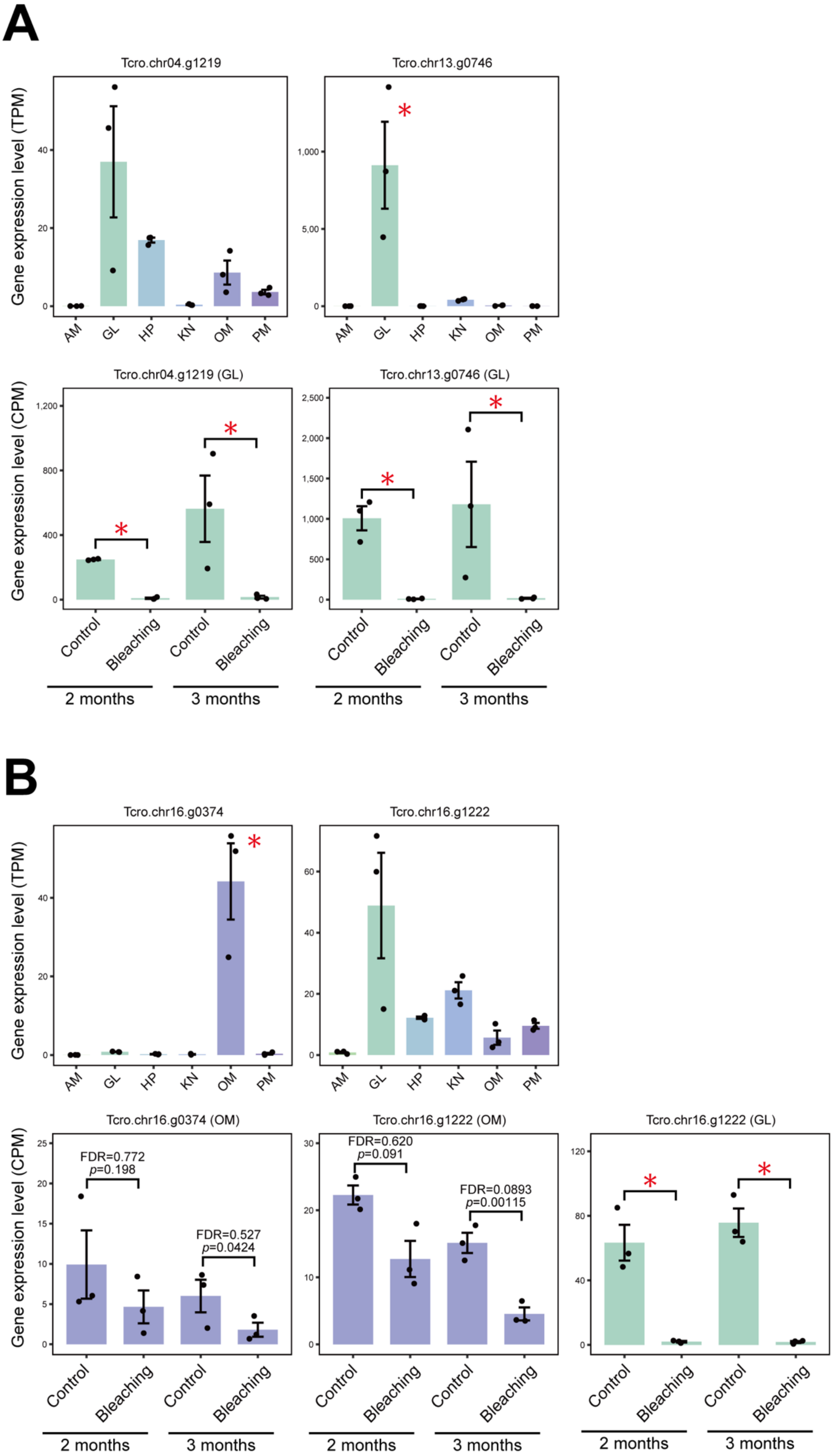
A
Tcro.chr04.g1219
Tcro.chr13.g0746
Gene expression level (TPM)
AM
GL
HP
KN
OM
PM
Tcro.chr04.g1219 (GL)
Tcro.chr13.g0746 (GL)
Gene expression level (CPM)
Control
Bleaching
2 months
3 months
B
Tcro.chr16.g0374
Tcro.chr16.g1222
Tcro.chr16.g0374 (OM)
Tcro.chr16.g1222 (OM)
Tcro.chr16.g1222 (GL)
FDR=0.772
p=0.198
FDR=0.527
p=0.0424
FDR=0.620
p=0.091
FDR=0.0893
p=0.00115

**Fig. S10.**

Gene expression of SLC4 and SLC 26 with characteristic expression patterns. (**A**) Tissue-specific gene expression. (**B**) Dark-induced bleaching experiment. * Tissue-specific HEGs or DEGs.

**Legends for Supplementary Tables**

**Table S1.**

Genome assembly and gene prediction statistics. BUSCO completeness of the genome and proteome was calculated using compleasm v.0.2.6 and BUSCO v.5.4.7, respectively. S, single-copy BUSCO; D, duplicated BUSCO. Telomere repeats (AACCCT)n were counted with tidk. * RefSeq representative genome (as of 2025/10/1). ** Li, J. Chromosome-level genome assembly and annotation of rare and endangered tropical bivalve, *Tridacna crocea*. figshare. Dataset. https://doi.org/10.6084/m9.figshare.24264646 *** Li *et al.* (*18*)

**Table S2.**

BLASTP top hits of *T. crocea* gene models against the SwissProt database. * Based on NCNI taxonomy (as of 2024/5). ** Gene prediction was performed in this study. *** The longest transcript variant for each locus is selected.

**Table S3.**

Gene models used for analysis by Orthofinder.

**Table S4.**

Fossil calibration points used for divergence time estimation.

**Table S5.**

RNA-seq data used for gene prediction.

**Table S6.**

List of tissue-specific HEGs.

**Table S7.**

Functional enrichment analysis for adductor muscle (AM)-specific HEGs.

**Table S8.**

Functional enrichment analysis for gill (GL)-specific HEGs.

**Table S9.**

Functional enrichment analysis for hepatopancreas (HP)-specific HEGs.

**Table S10.**
Functional enrichment analysis for kidney (KN)-specific HEGs.

**Table S11.**
Functional enrichment analysis for outer mantle (OM)-specific HEGs.

**Table S12.**
Functional enrichment analysis for pedal mantle (PM)-specific HEGs.

**Table S13.**
List of DEGs in the dark-induced bleaching experiment.

**Table S14.**
Functional enrichment analysis for DEGs in the gill (2 months, downregulated).

**Table S15.**
Functional enrichment analysis for DEGs in the gill (3 months, downregulated).

**Table S16.**
Functional enrichment analysis for DEGs in the outer mantle (2 months, downregulated).

**Table S17.**
Functional enrichment analysis for DEGs in the outer mantle (3 months, downregulated).

**Table S18.**
Publicly available *T. squamosa* RNA-seq data used for de novo transcriptome assembly.
